# Glioblastoma Treatment by Systemic Actinium-225 α-particle Dendrimer-radioconjugates is Improved by Chemotherapy

**DOI:** 10.1101/2024.10.17.618960

**Authors:** Rajiv Ranjit Nair, Aira Sarkar, Pooja Hariharan, Kathleen L. Gabrielson, Tony Wu, Chang Liu, Anjalia Santosh, Wathsala G.H.M Liyanage, Zaver M Bhujwalla, Marie-France Penet Vidaver, Rangaramanujam M Kannan, Stavroula Sofou

## Abstract

**RATIONALE:** The poor prognosis of glioblastoma is largely due to drug resistance and tumor location that, together, make it difficult to treat aggressively without affecting the rest of the brain.

**METHODOLOGY:** High-energy, short-range (40-80µm) dendrimer-delivered α-particles could address both challenges, because (1) they cause complex, highly cytotoxic double-strand DNA breaks, and (2) irradiation of the neighboring brain is minimal, since dendrimers selectively delivers them to tumors. Since cancer cells that are not directly hit by α-particles will likely not be killed, the patterns of tumor irradiation affect efficacy. Systemically injected dendrimers extensively accumulate in glioblastomas, where they are taken up by tumor associated macrophages (TAMs), which tend to infiltrate tumors. We hypothesized that dendrimers labeled with α-particle emitters, when being carried by TAMs, could more evenly irradiate glioblastomas, improving survival. In this study, the efficacy of dendrimers radiolabeled with the α-particle emitter actinium-225 (dendrimer-radioconjugates) was evaluated when administered alone and/or after temozolomide, in a syngeneic immune-competent orthotopic GL261-C57BL/6 mouse model.

**RESULTS:** Systemically-administered dendrimer-radioconjugates, at activities that did not result in long-term toxicities, prolonged survival of mice with orthotopic GL261 tumors, compared to standard-of-care temozolomide (39 vs 31 days mean survival, p=0.0061) and non-treated animals (30 days, p=0.0009). Importantly, injection of temozolomide 24 hours before administration of dendrimer-radioconjugates *further improved survival remarkably* (44 days). This improvement in efficacy was attributed to: (1) the significant increase (by 33%) in tumor absorbed doses delivered by dendrimer-radioconjugates when injected after chemotherapy, without altering normal organ dosimetry, while sparing the tumor-surrounding healthy brain; (2) the potentially deeper tumor penetration of dendrimer-radioconjugates, suggested by the enhancement of dendrimer penetration within GL261-spheroids, employed as model tumor-avascular regions and/or TAM-free regions; and/or (3) the formation of a more lethal cocktail when both modalities acted on same cancer cells, that was correlated with increased levels of dendrimer-radioconjugates associating with GL261 cells *in vitro* and with greater incidences of karyomegaly *in vivo*.

**CONCLUSIONS:** This study demonstrates the potential of a ‘brain tumor targeted’ systemic actinium-225 radiopharmaceutical therapy that inhibits growth of glioblastoma cells and prolongs survival of mice with orthotopic brain tumors, further improved by standard-of-care temozolomide, without notable toxicities.

## Introduction

Glioblastoma is an aggressive disease with a tragically short survival of a few months after diagnosis, despite treatment with multi-modal therapies [1, 2]. The poor prognosis is largely due to the tumor location, that makes it difficult to treat aggressively without affecting the rest of the brain, as well as due to the development of tumor resistance [3]. Furthermore, the relatively limited permeability of the blood brain tumor barrier (BBTB) may limit the extent of therapeutics’ accumulation in the tumor(s) [4].

Alpha-particle radiopharmaceutical therapy (αRPT) is a promising modality for such difficult to treat tumors. High-energy, short-range (40-80µm in tissue) α-particles unequivocally kill cancer cells by causing complex double-strand DNA-breaks [5]. αRPT is essentially impervious to resistance [5–7], if optimally delivered, and is ideal for treating glioblastoma, because irradiation of the nearby brain could be minimal. However, cancer cells not being hit by α-particles will likely not be killed [8]. Previous clinical and/or preclinical αRPT approaches for glioblastoma have employed vectors targeting the tumor vasculature and/or markers on cancer cells [9, 10]. Although these approaches demonstrated promise, they have failed to elicit long lasting responses; this could be partly due to inadequate delivery resulting in spatially non-uniform irradiation of tumors, which is the root cause of αRPT treatment failure in the clinic [11]. An approach that delivers αRPT to tumor regions uniformly, may enhance the therapeutic outcomes [7, 8].

Hydroxyl poly(amidoamine) dendrimers, are tree-like spherical ultrasmall nanoparticles [12], which have been previously demonstrated to: (1) selectively and extensively associate with microglia/activated macrophages; (2) selectively extravasate from circulation into intracranial gliomas by crossing the BBTB but not crossing the blood brain barrier (BBB) of the healthy brain; (3) be extensively distributed within brain tumors by being associated with tumor-associated macrophages (TAMs) [13]; and (4) clear fast from off-target organs renally, even in humans [13–16]. The hydroxyl PAMAM dendrimer platform has been shown to be safe in humans, with positive Phase 2 results in microphage/inflammation-targeted systemic treatments for hospitalized severe COVID-19 patients.

Given that glioblastomas are characterized by high levels of TAMs [17], to enable selective tumor uptake and more uniform patterns of tumor irradiation by αRPT, we engineered a delivery strategy where αRPT was attached on systemically injected dendrimers. Unlike other reports, our aim was neither to deplete nor to reprogram TAMs [18], but instead to employ TAMs as carriers of the systemically injected αRPT, and to help infiltrate and more uniformly irradiate glioblastoma tumors with α-particles.

In this study, Actinium-225 ([^225^Ac]Ac) dendrimer-radioconjugates were characterized *in vitro* using murine GL261 glioblastoma cells and macrophages, and *in vivo* on C57BL/6 immune-competent mice with orthotopic GL261 tumors, and their efficacy to prolong survival was compared to that of the standard-of-care temozolomide. The dosimetry and long-term toxicities were also evaluated.

## Materials and Methods

Reagent sourcing is detailed in SI. The actinium-225 used in this research was supplied by the U.S. Department of Energy Isotope Program managed by the Office of Isotope R&D and Production. Indium-111 (^111^In), indium chloride, was purchased from BWXT (Ontario, Canada).

### Dendrimer characterization and (radio)labeling

Hydroxyl-terminated Generation-6 poly(amidoamine) dendrimers were conjugated with chelators DOTA (DOTA-dendrimer) or DTPA (DTPA-dendrimer) [19] and/or were fluorescently labeled (Cy5-dendrimers), as preciously described [13]. Size and zeta potential were measured by a Nanoseries Zetasizer (Malvern Instruments Ltd, Worcestershire, UK).

To radiolabel dendrimers with [^225^Ac]Ac (or [^111^In]In), DOTA-dendrimer (or DTPA-dendrimer) was processed following a one-step radiolabeling protocol, and measurement of specific activity and stability of radiolabeling were evaluated as previously described [7, 8] and also detailed in SI.

### Cell lines

The murine glioblastoma cell line, GL261, was obtained from DCTD /Tumor Repository (National Cancer Institute, Frederick, Maryland). Murine BV2 macrophages were obtained from the Children’s Hospital of Michigan Cell Culture Facility. Cells were cultured using Roswell Park Memorial Institute (RPMI) and Dulbecco’s modified Eagle’s media (DMEM), respectively, each being supplemented with 10% FBS, 100units/mL Penicillin and 100µg/mL Streptomycin in an incubator at 37°C and 5% CO_2_. For M1 activation, macrophages were incubated in DMEM supplemented with 5% FBS, 100units/mL Penicillin and 100µg/mL Streptomycin, containing 100ng/mL of Lipo-polysaccharide, at 37°C and 5% CO_2_ for 48 hours [20]. For M2 activation, macrophages were incubated in DMEM media supplemented with 10% FBS, 100units/mL Penicillin and 100µg/mL Streptomycin, and 8ng/mL of murine IL-4, in an incubator at 37 °C and 5% CO_2_ for 48 hours [21].

### IC_50_ of Temozolomide

Cell survival was measured after two cell-doubling times (192 hours for GL261; 70 hours for activated BV2) following completion of a 6-hour incubation with TMZ, of cells on 96-well plates (20,000 cells per well), using a 3-(4,5-dimethylthiazol-2-yl)-2,5-diphenyltetrazolium bromide (MTT) assay.

### Clonogenic cell survival assay and measurement of radioactivity associated per cell

Clonogenic cell survival fractions were measured after exposure of cells for 6 hours to actinium-225 and/or TMZ, and then plating in tissue culture dishes till the onset of colony formation, as previously described [7, 8] and detailed in SI.

To measure the mean radioactivity associated per cell, for the highest two activity concentrations, post incubation, on a fraction of the cell suspension, the γ-photon emissions of bismuth-213 were measured (at secular equilibrium), and the total activity was then divided by the total number of live cells, which were counted using Trypan Blue.

### Spheroids

Spheroids were formed by plating 1,000 GL261 cells per well onto ultra-low adhesion U-shaped 96-well plates (in 1.8% v/v Matrigel^TM^) following centrifugation at 1,023RCF for 10 minutes. Spheroids were then let grow until they reached 400μm diameter.

To evaluate the spatiotemporal microdistributions of dendrimers (and/or of free TMZ-surrogate) within GL261 spheroids, the latter were incubated with fluorescent Cy5-dendrimers (ex/em: 651/670 nm) (and/or CFDA-SE, employed as a fluorescent surrogate of TMZ; ex/em: 492/517 nm) at a final concentration of 10μg/mL (and/or 10μM). During incubation (“uptake phase”), spheroids were sampled at different time points and were then transferred to fresh media and were sampled during the “clearance” phase (of agents from spheroids). Sampled spheroids were frozen, sectioned and imaged, and the radial distributions of each fluorescent species were quantified using an inhouse-eroding code as previously described [7, 8] and detailed in SI.

To evaluate the effect of treatments on the extent of regrowth, post completion of incubation with [^225^Ac]Ac-DOTA-dendrimers and/or with TMZ, spheroids were transferred to fresh media and their size was monitored daily, until the growth rate of non-treated spheroids reached an asymptote (∼10 days). At that point, each spheroid was plated on a separate well of a 96-flat well adherent cell culture plate and was allowed to grow till the cells from the non-treatment group reached 70− 80% confluency. All cells were then trypsinized, and cell counts were normalized to the number of the non–treated cells to obtain a “percent spheroid regrowth relative to no treatment”.

### Animal study

Mice were housed in filter top cages with sterile food and water, and all studies were performed in compliance with the Institutional Animal Care and Use Committee protocol guidelines. Four to six-week, 20g C57BL/6J male mice (Jackson Laboratory, Bar Harbor, ME, USA), were intracranially inoculated with 100,000 GL261 cells in 2µL as previously described [22]. To confirm the presence of tumor, before randomly assigning mice to different groups, mouse brain images were acquired on a multi-nuclear BioSpec 70/30 PET-MR 7T scanner (Bruker Biospin MRI Inc., Billerica, MA, USA) using a T2-weighted fast spin-echo rapid imaging with refocused echo (RARE) sequence, as described in the SI.

In treatment studies, when indicated, TMZ (100µL of 80mg TMZ/kg mouse) was administered intraperitoneally 24 hours before intravenous injection of 100µL [^225^Ac]Ac-DOTA-dendrimers. When animal weight loss was ≥25%, following euthanasia, the tumor and normal organs were harvested, fixed and stained with H&E, and were evaluated for pathological findings. The highest mass of injected dendrimer (1.5mg/Kg mouse) was significantly lower (by two orders of magnitude) than dendrimer concentrations used previously without causing side effects [14].

Biodistributions were evaluated by intravenously administering 100µL of 740kBq [^111^In]In-DTPA-dendrimers. At different time points, animals were sacrificed, and tumor and normal-organs were excised, weighted, and a γ-counter was used to measure their associated activity, which was then decay-corrected and reported as percentage of the initially injected activity per gram of tissue (%IA/g).

### Dosimetry

The biodistributions of [^111^In]In-DTPA-dendrimers were utilized to calculate the absorbed doses for [^225^Ac]Ac-DOTA-dendrimers, as previously described [23], using the software package RAPID Dosimetry (Baltimore, MD).

### Statistical Analysis

Results were reported as the arithmetic mean of n independent measurements ± the standard deviation. One-way ANOVA and/or the unpaired Student’s t test were used to calculate differences between experimental groups. Comparison of survival on Kaplan-Meier plots was performed using the log-rank test. P-values <0.05 were considered significant. * indicates 0.01<p-values<0.05; **<0.01, ***<0.001.

## Results

### Dendrimer radiolabeling

Dendrimers were stably radiolabeled with [^225^Ac]Ac (and/or with [^111^In]In, respectively), resulting in reasonable levels of specific activities (Table 1). The 7.6±0.6nm (PDI = 0.05±0.02) dendrimers had close to neutral zeta potential (Table 2) both in pH 7.4 and 6.2 (chosen to represent the acidic interstitium in spheroids [24], Figure S1).

**TABLE 1.**
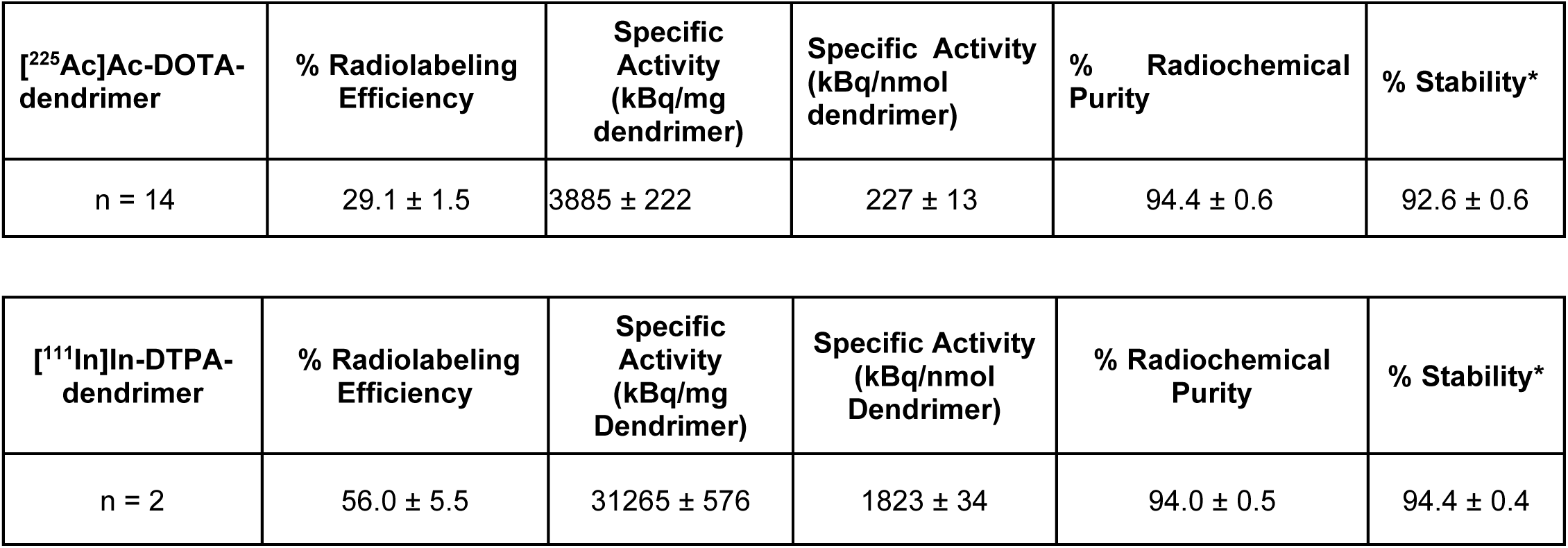
Characterization of radiolabeled dendrimers. *Stability assessed in media at pH 7.4 and 37°C after 24 hours of incubation. Shown are the mean values ± the standard deviation of n independent measurements.

| [$^{225}\text{Ac}$]Ac-DOTA-dendrimer | % Radiolabeling Efficiency | Specific Activity (kBq/mg dendrimer) | Specific Activity (kBq/nmol dendrimer) | % Radiochemical Purity | % Stability* |
| --- | --- | --- | --- | --- | --- |
| n = 14 | $29.1 \pm 1.5$ | $3885 \pm 222$ | $227 \pm 13$ | $94.4 \pm 0.6$ | $92.6 \pm 0.6$ |

| [$^{111}\text{In}$]In-DTPA-dendrimer | % Radiolabeling Efficiency | Specific Activity (kBq/mg Dendrimer) | Specific Activity (kBq/nmol Dendrimer) | % Radiochemical Purity | % Stability* |
| --- | --- | --- | --- | --- | --- |
| n = 2 | $56.0 \pm 5.5$ | $31265 \pm 576$ | $1823 \pm 34$ | $94.0 \pm 0.5$ | $94.4 \pm 0.4$ |

**TABLE 2.** Zeta potential of dendrimers functionalized with DOTA and/or DTPA. Shown are the mean values ± the standard deviation of n = 3 independent measurements.

| Zeta potential (mV) | DOTA-dendrimer | Cy5-dendrimer |
| --- | --- | --- |
| pH 6.2 | $-0.17 \pm 0.10$ | $-0.13 \pm 0.12$ |
| pH 7.4 | $0.15 \pm 0.26$ | $0.21 \pm 0.30$ |

### Macrophages were less sensitive to actinium-225 than GL261 glioblastoma cells in vitro

Figure 1A shows that GL261 murine glioblastoma cells were more sensitive to α-particle radiation, in the form of [^225^Ac]Ac-DOTA, compared to resting and/or activated BV2 macrophages (*p*-value <0.01). [^225^Ac]Ac-DOTA was not expected to significantly and/or preferentially associate with any cell type [19], enabling for relatively uniform and comparable irradiation of all cell types. In Figure 1B, upon exposure to [^225^Ac]Ac-DOTA-dendrimers, glioblastoma cells again exhibited the greatest sensitivity. However, because of the significant association of dendrimers with activated macrophages (Figure 1C), the latter exhibited lower survival factions upon exposure to [^225^Ac]Ac-DOTA-dendrimers than to [^225^Ac]Ac-DOTA.

**FIGURE 1.**
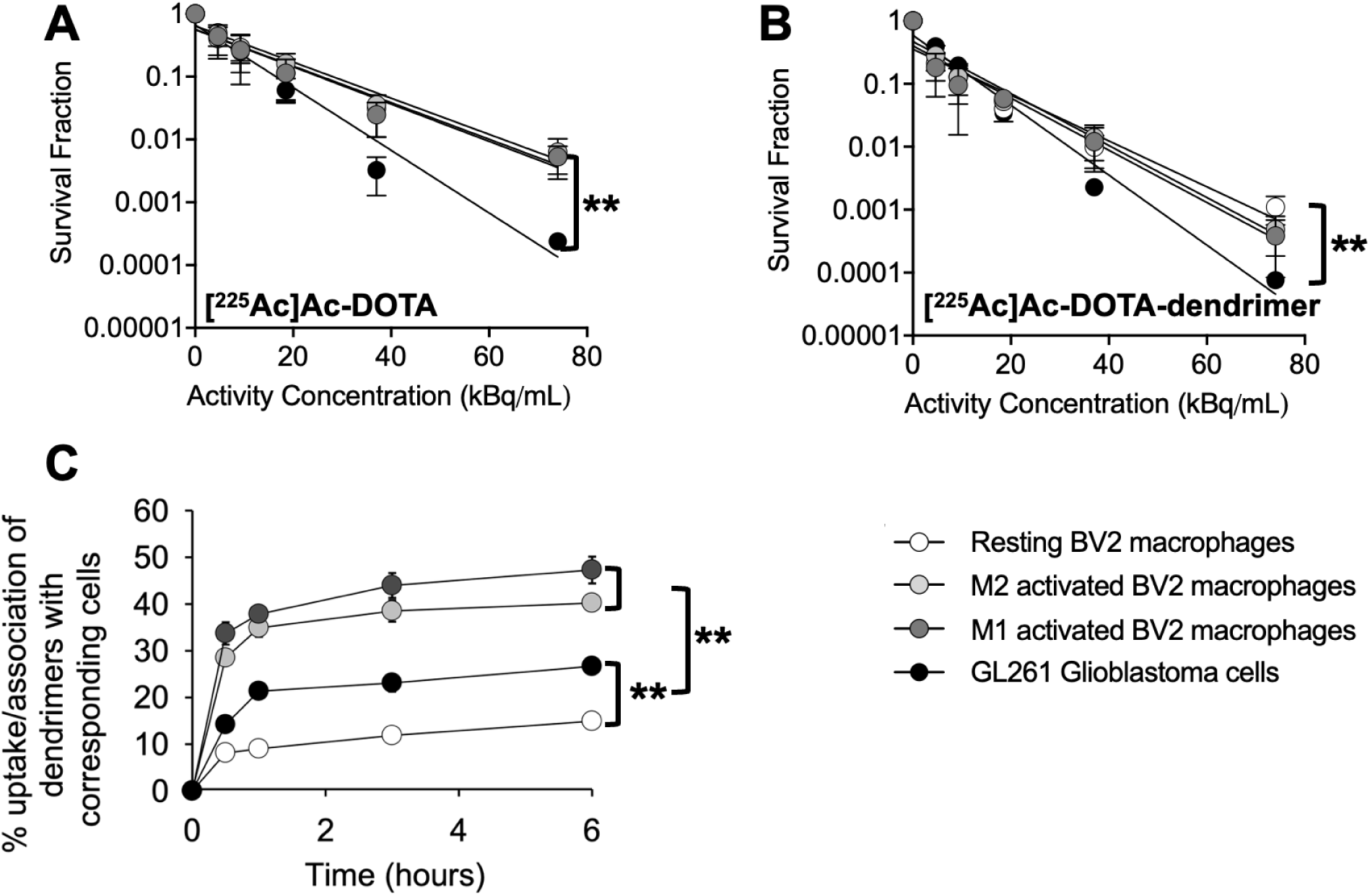
Clonogenic cell survival of GL261 glioblastoma cells and BV2 macrophages exposed to (A) [^225^Ac]Ac-DOTA and/or (B) [^225^Ac]Ac-DOTA-dendrimers, for 6 hours at 37°C. The concentration of dendrimer-radioconjugates in (B) was kept constant at all activity concentrations by adjusting with cold dendrimer. (C) Time-dependent extent of uptake/association of fluorescently-labeled Cy5-dendrimers with resting and activated macrophages, and/or GL261 glioblastoma cells. Uptake Method in C: Cells in suspension (2×10^6^ cells/mL) were incubated with Cy5-dendrimers (20.4μg/mL) at dendrimer-to-cell ratio equal to 50×10^6^-to-1. Mean values ± the standard deviation of n=3 independent runs are shown. ** indicates *p*-value <0.01.

### Exposure to Temozolomide increased association of (radiolabeled) dendrimers with cells enhancing cytotoxicity

Given that TMZ is a standard-of-care chemotherapeutic [25], the effect of actinium-225 on GL261 glioblastoma cells (and on macrophages) was also evaluated in the presence of TMZ. Figures 2A and 2B (GL261 and macrophages, respectively) show that the extents of clonogenic cell survival correlated strongly with the activities associated per cell. Additionally, increasing concentrations of TMZ in the incubation media (gray and black fills on same symbols, at concentrations lower than the IC_50_, Figure S2), increased the activity levels associated per cell resulting in even lower survival fractions. This finding was independent of the form of actinium-225 ([^225^Ac]Ac-DOTA, shown in circles, and/or [^225^Ac]Ac-DOTA-dendrimers, in squares) and/or of the activity concentration during incubation (top/lower set of symbols corresponds to 37/ 74 kBq/mL, from the clonogenic cell survival vs activity plots shown in Figure S3). The increased uptake of Cy5-dendrimers by GL261 cells and/or macrophages in the presence of TMZ (Figures 2C and 2D) was also observed in the absence of radioactivity and was more pronounced when the exposure to TMZ preceded the introduction of dendrimers into the incubation medium.

**FIGURE 2.**
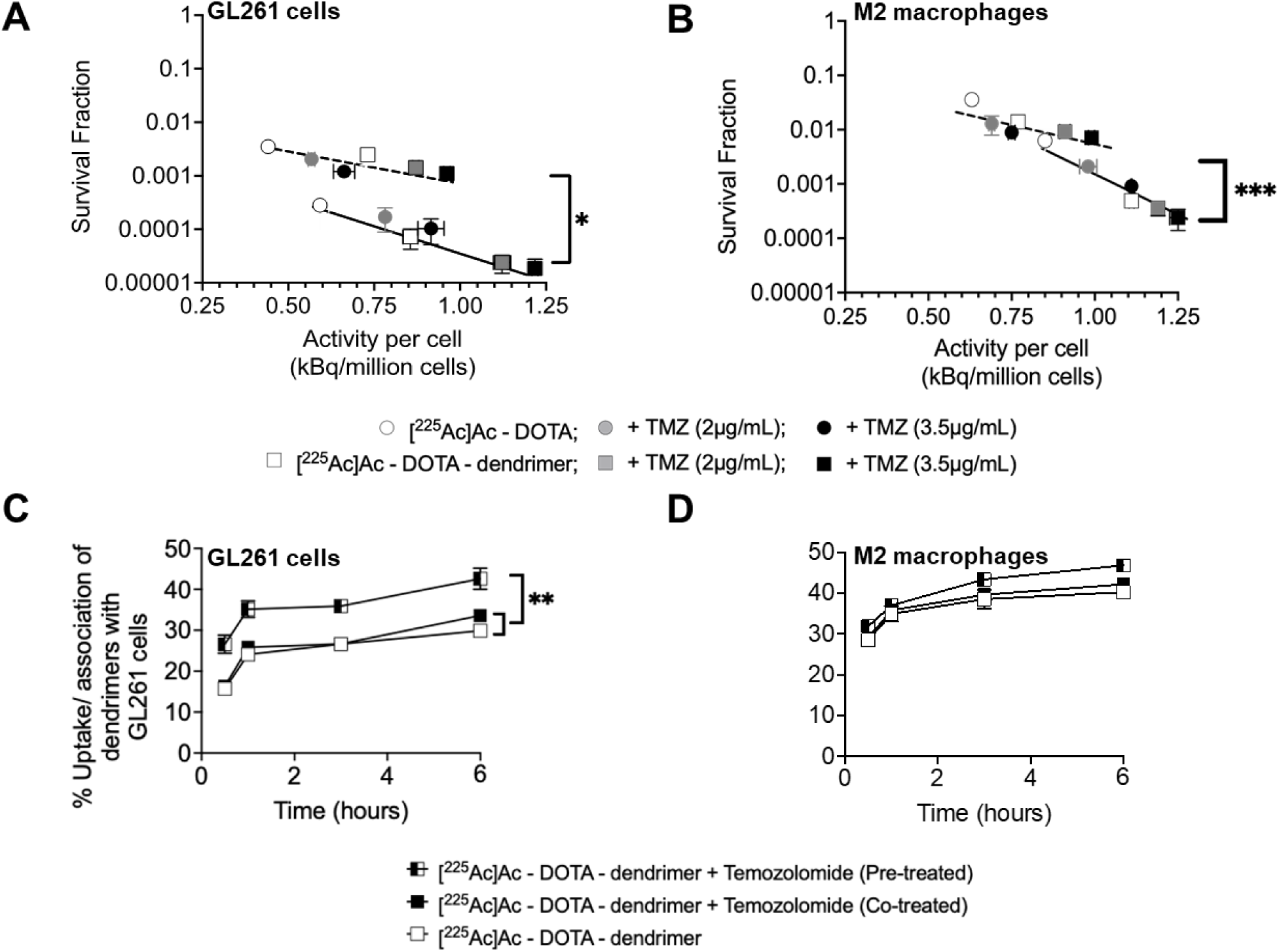
Top panels: Clonogenic cell survival of (A) GL261 glioblastoma cells and (B) BV2 M2 macrophages, exposed to [^225^Ac]Ac-DOTA (circles) and/or [^225^Ac]Ac-DOTA-dendrimers (squares), in the presence (gray/black) and/or absence (white) of TMZ, as a function of the activity associated per cell, after a 6 hour incubation ([dendrimer]=10µg/mL). Mean values ± standard deviation of n=2 independent runs are shown. The dotted (and solid) line is best fit of activity per cell for cells incubated with 37 (and 74) kBq/mL, respectively.

Low panels: Time-dependent uptake/association of Cy5-dendrimers with (C) GL261 glioblastoma cells and (D) M2 macrophages that were either pretreated with TMZ (half-black/half-white symbols) or concurrently exposed to both TMZ and the dendrimers (black symbols), compared to the cell uptake of dendrimers in the absence of TMZ in the incubation medium (white symbols). Cells in suspension (2×10^6^ cells/mL) were incubated with dendrimers (20.4µg/mL) at dendrimer-to-cell ratio equal to 50×10^6^-to-1. TMZ concentration was 18.03 µM. Mean values ± standard deviation of n = 3 independent runs are shown. * indicates 0.01<*p*-value<0.05; **<0.01.

### GL261 spheroid studies

The time-integrated microdistributions of Cy5-dendrimers within GL261 spheroids - which were utilized as surrogates of the tumors’ avascular regions - were heterogeneous (Figure 3A), partly because of their association with cancer cells [26] and the absence of infiltrating macrophages. Contrary, the small fluorescent molecule CFDA-SE, used as surrogate of TMZ, exhibited significant penetration within the spheroids (Figure 3B) (spatial profiles of each species vs time are shown in Figure S4). The Cy5-dendrimers, however, exhibited deeper penetration (Figure 3C; the distribution at mid height was shifted towards the core by 12 µm) when GL261 spheroids were pre-incubated with TMZ, possibly due to partial killing of some of GL261 cells.

**FIGURE 3.**
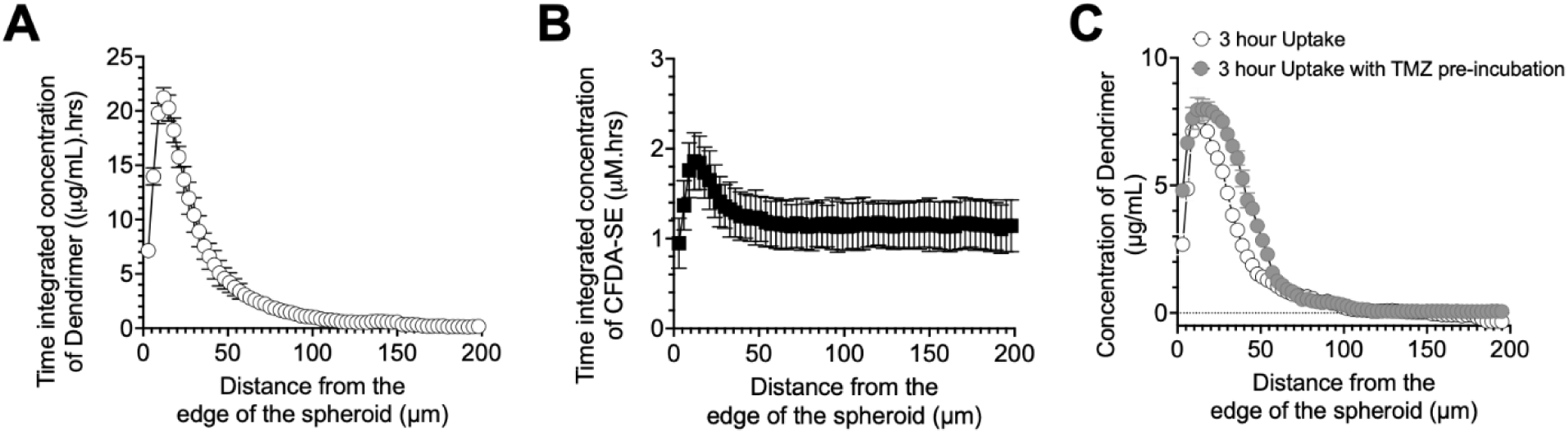
Time integrated radial microdistributions in 400µm-in-diameter GL261 spheroids of (A) Cy5-labeled dendrimers, and (B) CFDA-SE, utilized as a temozolomide (TMZ) fluorescent surrogate. Spatial profiles of each agent vs time are shown in Figure S4. (C) Pre-treatment of spheroids with TMZ improved the penetration of Cy5-dendrimers. Data points indicate the mean values ± the standard deviations of n=3 different spheroids per time point.

The percent inhibition of spheroid regrowth following exposure to both agents (TMZ and dendrimer-radioconjugates, gray bars) was greater (55.95%, 71.05% and 82.22% at 3, 6 and 9kBq/mL, respectively) than the added regrowth inhibition following exposure to either agent alone (Figure 4, white bars and TMZ-labelled black bar; 45.95%, 66.33% and 72.39%, respectively). On cell monolayers, improved killing was observed when both agents acted on same cells. In spheroids, even under the non-uniform α-particle spatiotemporal microdistributions (suggested by the limited dendrimer penetration shown in Figure 3A), a significant fraction of the cancer cells, notably those located within the first at least 50 µm from the spheroid edge, was exposed to both modalities, possibly contributing to the observed enhancement in regrowth inhibition.

**FIGURE 4.**
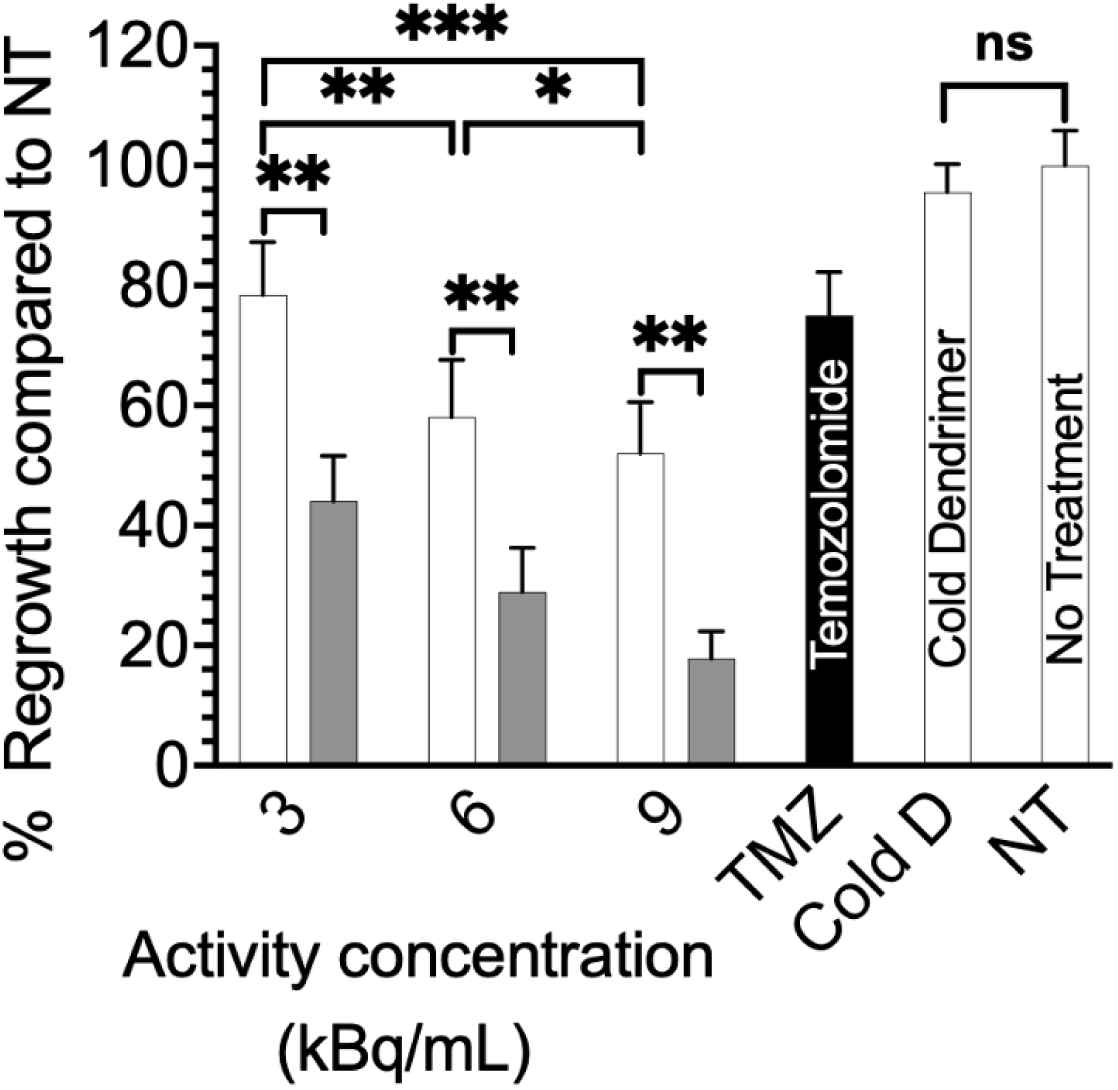
The extent of inhibiting regrowth (used as indirect surrogate of tumor recurrence) of 400µm-in-diameter GL261 spheroids treated with [^225^Ac]Ac-DOTA-dendrimer (white bars), temozolomide (black bar) and/or their combination (gray bars) relative to untreated spheroids and the cold dendrimer. Spheroids were exposed to dendrimer-radioconjugates for 2 hours in the absence or presence of TMZ (pre-treated for 22 hours), and/or to TMZ alone (24 hours). Mean values ± the standard deviations of n=6 spheroids per condition (n=3 independent dendrimer-radioconjugate preparations) as shown. *** indicates *p*-value <0.001; **<0.01; *<0.05.

### In vivo studies

### Biodistributions and dosimetry: pre-administration of TMZ selectively increased the tumor uptake of systemically injected dendrimer-radioconjugates

The biodistributions of [^111^In]In-DTPA-dendrimer, alone and/or following the injection of TMZ, 24 hours earlier (Figures S5 and S6, and Tables S1 and S2), were employed to calculate the dosimetry of [^225^Ac]Ac-DOTA-dendrimer, since [^111^In]In has been previously validated as a surrogate of the parent actinium-225 [27–29]. All daughters’ (α-particle and electron) energy delivered to tumors by dendrimers was assumed to be absorbed by the tumor, because the activity associated with and retained by the tumors was attributed primarily to the dendrimer-radioconjugates that were internalized by TAMs (Figure S7 shows that macrophages internalized almost 30% of the dendrimers to which they were exposed). Additionally, the unusually long clearance kinetics of dendrimer-radioconjugates from tumors (the clearance half-life after tumor’s maximum uptake was: 72 hours) [30], that was even longer than the clearance kinetics of internalizing-antibodies (approximately 24-36 hours clearance half-life after maximum tumor uptake) [27, 28], also supported this choice.

Interestingly, when TMZ was injected i.p. 24 hours before systemic injection of dendrimer-radioconjugates (Figure 5, Table 3, Table S3), only the tumor absorbed dose significantly increased (from 0.24±0.01 to 0.32±0.01Gy; *p*-value=0.0006). Notably, the off-target organ uptake of dendrimer-radioconjugates was not affected by pre-injecting TMZ.

**FIGURE 5.**
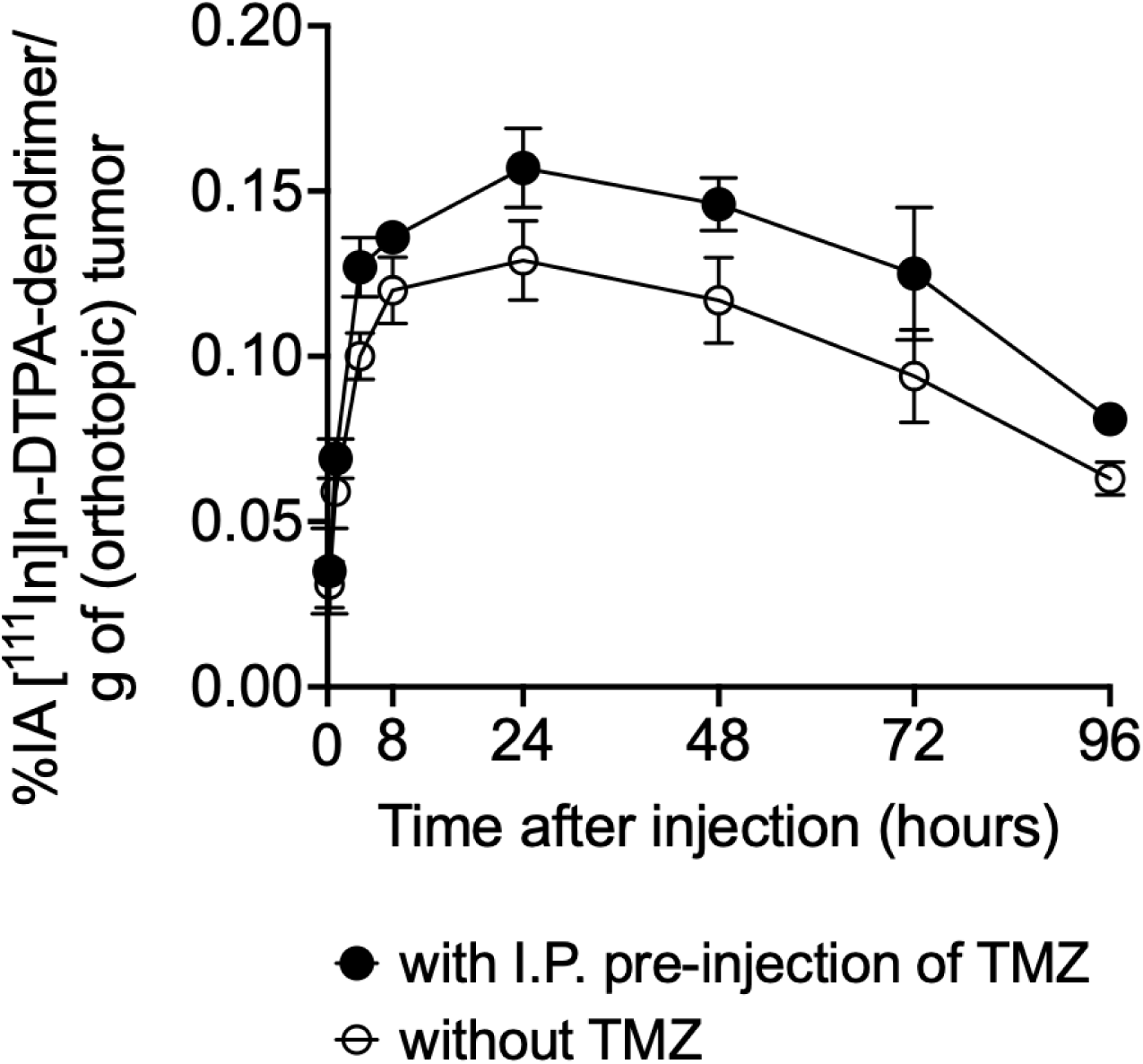
The uptake of systemically injected [^111^In]In-DTPA-dendrimer by orthotopic GL261 glioblastoma tumors in C57BL/6 mice, significantly increased when TMZ (80mg/Kg mouse) was intraperitoneally injected 24 hours before I.V. administration of dendrimer-radioconjugates. At different timepoints animals were sacrificed and the tumor (and tissues) were dissected, weighted, and the associated activity was measured using a gamma-counter, as previously reported [27–29]. The decay-corrected % injected activity per mass of tumor (for n=3 animals per time point) is shown. Notably, normal brain’s dendrimer-radioconjugate uptake was not affected by TMZ pre-injection (Table 3, Figures S5, S6).

**TABLE 3:** Dosimetry for systemically injected [^111^In]In-DTPA-dendrimer in (A) the absence of Temozolomide (TMZ), and (B) following intraperitoneal injection of TMZ, 80mg/Kg mouse, 24 hours earlier. Calculated for 29.6kBq [^225^Ac]Ac-DOTA-dendrimer per 20g mouse using RAPID Dosimetry (Baltimore, MD) (see Table S3 for 22.2kBq/20g mouse). Reported are the mean values and standard deviations of dose evaluated for n=3 mice per time point. ***p-value<0.001.

| Organ | Absorbed Dose (Gy) |  |
| --- | --- | --- |
|  | Without TMZ | With I.P. pre-injection of TMZ |
| Tumor | 0.2 ± 0.0*** | 0.3 ± 0.0*** |
| Heart | 0.9 ± 0.0 | 0.9 ± 0.0 |
| Lungs | 0.0 ± 0.0 | 0.0 ± 0.0 |
| Liver | 0.2 ± 0.0 | 0.2 ± 0.0 |
| Spleen | 0.0 ± 0.0 | 0.0 ± 0.0 |
| Kidneys | 2.1 ± 0.1 | 1.9 ± 0.1 |
| Brain | 0.1 ± 0.0 | 0.1 ± 0.0 |
| Intestine | 0.0 ± 0.0 | 0.0 ± 0.0 |
| Stomach | 0.0 ± 0.0 | 0.0 ± 0.0 |

### Long-term toxicity studies

The MTD of [^225^Ac]Ac-DOTA-dendrimers (defined as the maximum activity at which no deaths were observed) was found to be greater than 29.6kBq and less than 44.4kBq per 20g tumor-free mouse. Long-term (11 months after administration of 29.6kBq [^225^Ac]Ac-DOTA-dendrimer) toxicities at the kidneys, heart and/or other normal organs were not observed by pathology examination (Figure 6A and Figure S8-(A)). However, at 44.4kBq injected activity, significant renal, splenic and hepatic toxicity was noted (Figure S8-(B)). The liver tissue exhibited significant hepatic lipidosis and the kidneys demonstrated tubular atrophy, vacuolization, karyomegaly and slight inflammation and fibrosis. The spleens exhibited iron accumulation, hemosiderin, indicative of past cellular death. Notably, the brain did not raise any long-term (11 months) pathological concerns: neither at 29.6kBq nor at 44.4 kBq (on those animals that survived).

**FIGURE 6.**
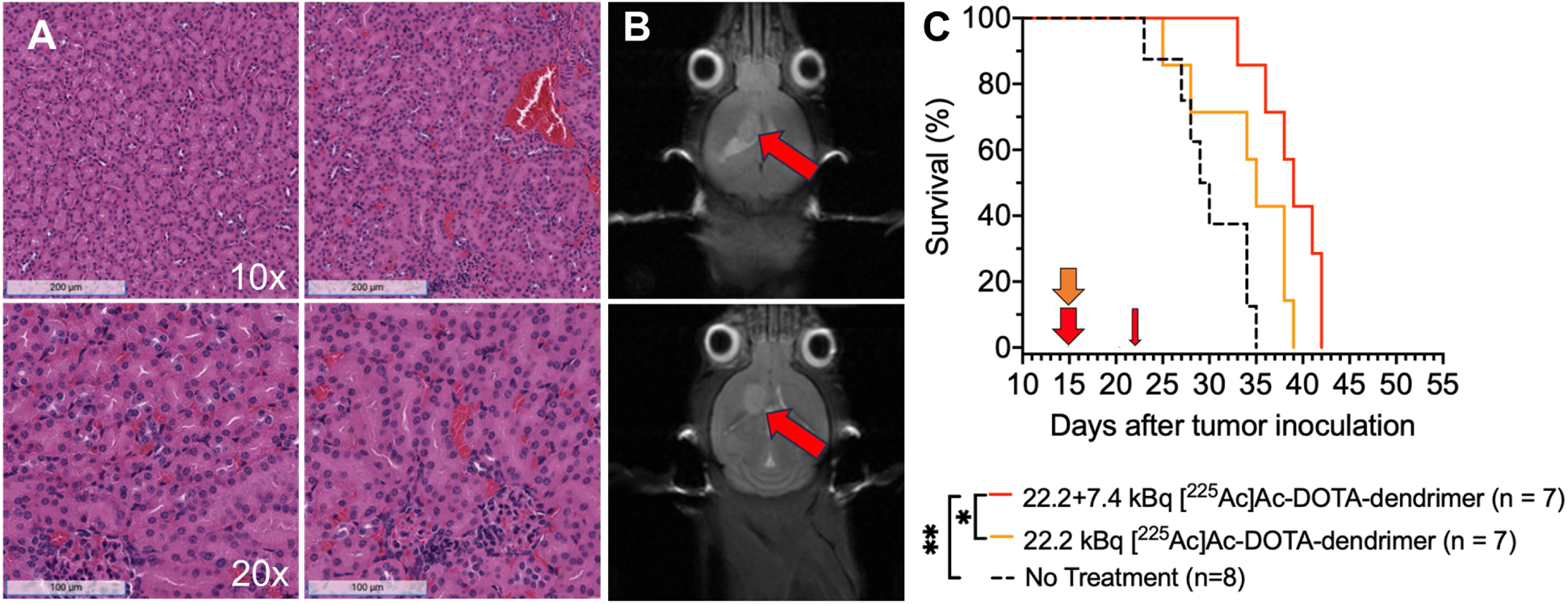
(A) Long-term renal toxicities (11 months after systemic administration of cumulative 29.6kBq [^225^Ac]Ac-DOTA-dendrimer, that clears via the kidneys) were not observed by pathology examination, on tumor-free C57BL/6 mice. (B) Characteristic MRI scans of C57BL/6 mice with orthotopic GL261 glioblastoma tumors (inoculated with 100,000 GL261 cells on day 0) with volumes averaging 9.7±5.7 mm^3^, a day after initiation of treatment. (C) Survival of same mouse model injected systemically with [^225^Ac]Ac-DOTA-dendrimer. Vertical arrows indicate treatment scheduling. Mice were treated with activities below the MTD (29.6kBq<MTD<44.4kBq /20g mouse). The study endpoint was determined by weight loss ≥25%. Statistical significance was evaluated by the log-rank test. ** indicates *p*-value <0.01; *<0.05.

### Response to treatment

Immune competent C57BL/6 mice with orthotopic murine GL261 glioblastoma tumors were treated when tumor volumes averaged 9.7±5.7mm^3^ (Figure 6B). Significantly improved survival, relative to no treatment (39 vs 30 days mean survival, p=0.0009), was observed for administered activities > 22.2 kB per 20g mouse (Figure 6C). As expected for αRPT [23], dose fractionation did not affect animal survival (Figure S9), although the time of first death was delayed when a higher fraction of activity was injected first. Additionally, the time of treatment initiation, day 10 (average tumor volume 1.5±0.7mm^3^) vs day 15, after tumor inoculation, did not affect the survival outcome (Figure S10 in supporting information).

Importantly, although administration of TMZ alone did not have an effect on animal survival relative to no treatment (gray line in Figure 7A, mean survival 31 days), when injected 24 hours before systemically injecting 29.6kBq [^225^Ac]Ac-DOTA-dendrimer *a significantly prolonged survival was observed* (red dashed line in Figure 7A, mean survival 44 days), which was even longer than the survival of mice that received radiotherapy alone (red solid line; mean survival 39 days, *p*-value=0.0017). On all treated groups, pathology evaluation of the tumor-brain interface did not reveal any adverse effects of αRPT on the neighboring healthy brain (Figure 7B). Pathology evaluation of the brain tumors when exposed to αRPT with pre-administered TMZ demonstrated increased population of karyomegalic tumor cells, in comparison to the chemotherapy-only cohort (Figure 7B, red framed regions).

**FIGURE 7.**
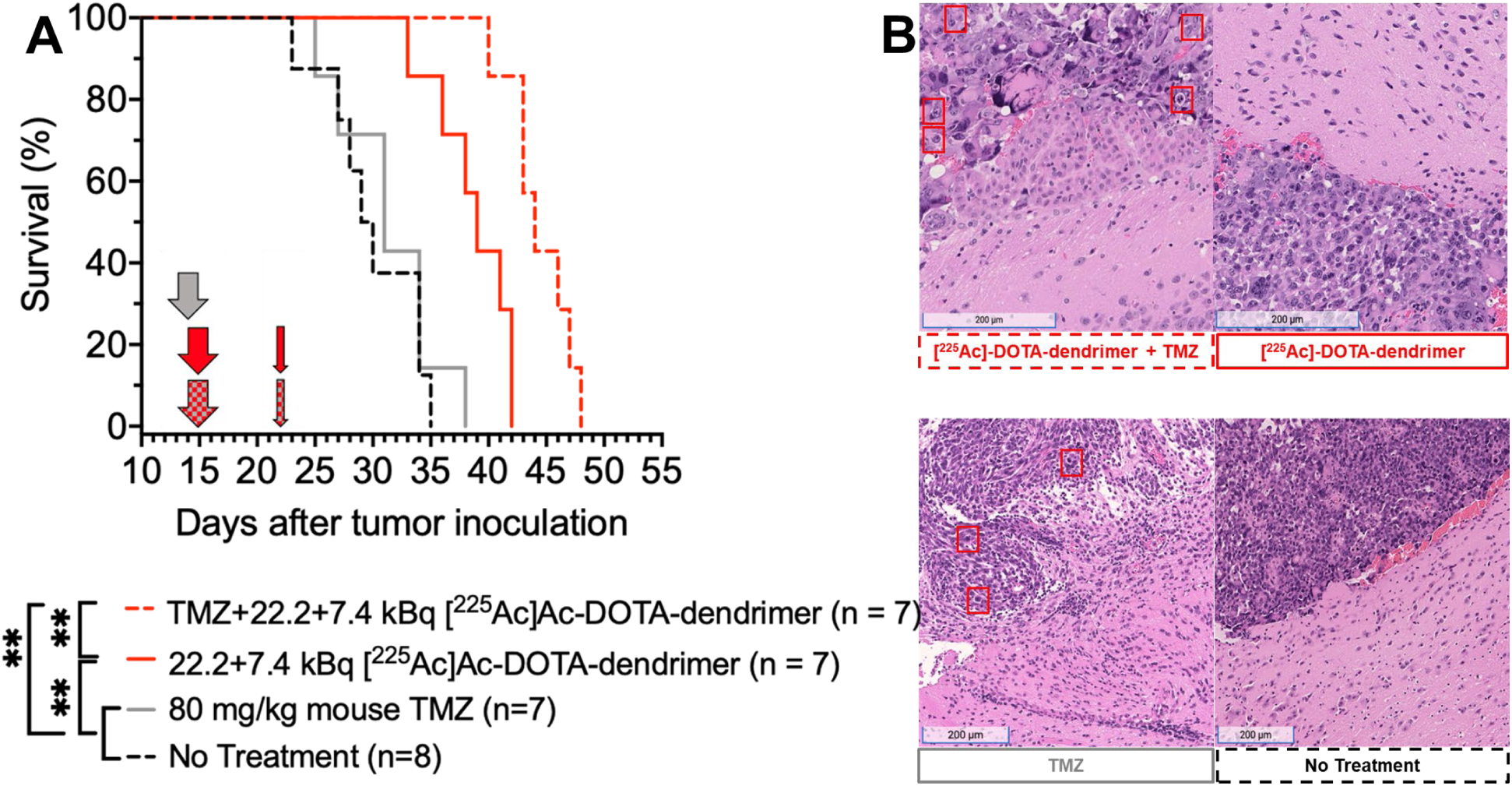
(A) Survival of C57BL/6 mice with orthotopic GL261 glioblastoma tumors treated with intravenous injection of [^225^Ac]Ac-DOTA-dendrimers (29.6kBq cumulative activity per 20g mouse, on day 15 and day 22 after tumor inoculation), with intraperitoneal injection of 80mg TMZ/Kg mouse (on day 14), and/or their combination relative to non-treated mice (TMZ on day 14, αRPT on day 15 and day 22). Vertical arrows indicate treatment scheduling. The study endpoint was determined by weight loss ≥25%. Statistical significance was evaluated by the log-rank test. ** indicates *p*-value <0.01. (B) Characteristic H&E-stained sections of the tumor-healthy brain interface.

In all treated groups, unless noted, no significant renal, hepatic or splenic toxicity was observed (Figure S11). In the chemotherapy-only cohort, mild hepatic lipidosis was observed. However, this was not exhibited by the cohort treated with the combination of αRPT and chemotherapy. All animal weights over time are reported in Figure S12.

## Discussion

On immune competent mice with syngeneic orthotopic GL261 glioblastoma, systemically injected α-particle dendrimer-radioconjugates — that selectively accumulated in brain tumors and, as previously shown, were selectively taken up by tumor-infiltrating TAMs [13] — significantly prolonged survival (39 days mean survival), compared to standard of care temozolomide (TMZ, 31 days) and to no treatment (30 days), at administered doses below the MTD. Interestingly, when 24 hours before injection of αRPT, a single dose of standard-of-care chemotherapy was administered - that itself did not affect survival – remarkable synergy in prolonging survival was observed (44 days). In all studies, therapy was initiated when tumor volumes were substantially grown (15 days after tumor inoculation). Notably, at the time of sacrifice, toxicities on the nearby brain were not detected in any treated group. [^225^Ac]Ac-labeled dendrimer-radioconjugates improved survival at tumor absorbed doses (0.2-0.3Gy) that were 2-3x lower than the tumor absorbed doses reported for vasculature-targeting [^225^Ac]Ac-labeled antibody-radioconjugates, which also improved survival [31], albeit on a different orthotopic glioblastoma mouse model. Although not directly comparable, the similar efficacy at dissimilar tumor absorbed doses could be partially explained by the different micro-patterns of tumor irradiation by α-particles between the two approaches: a more uniform, in the case of the lower absorbed doses delivered by dendrimers, vs. a more localized, higher dose irradiating mostly the vasculature.

We identified three reasons that partly explained the unexpected efficacy improvement when pre-injecting chemotherapy in this proof-of-concept study. First, at the whole-body scale, we observed increased tumor uptake of dendrimer-radioconjugates and, therefore, higher tumor delivered doses, when αRPT was administered 24 hours after chemotherapy. Our treatment scheduling was different from previous reports where αRPT was administered concurrently [31] with (higher doses of) chemotherapy, or where chemotherapy was administered after αRPT, which, however targeted the vasculature and was not intended to be distributed evenly within the tumor’s volume [32]. It is possible, that during the 24 hours that intervened, chemotherapy acted on the tumor vasculature [33] increasing its permeability. Second, pre-injection of chemotherapy could have “decompressed” the tumor microenvironment by killing some of the glioblastoma cells residing close to the tumor periphery [33]. Although we did not measure the spatial distributions of dendrimer-radioconjugates within glioblastoma tumors, our studies on evaluating the penetration profiles of fluorescently-labeled dendrimers in GL261 spheroids demonstrated deeper penetration when dendrimers were introduced after chemotherapy. Thirdly, we demonstrated that at the single-cell scale, when both αRPT and chemotherapy acted on same glioblastoma cells, the efficacy on these cells was higher than the efficacy of each agent alone, in agreement with previous reports [7, 34].

The normal brain exhibited low uptake of dendrimer-radioconjugates relative to brain tumors, and the tumor-neighboring brain was free of pathological findings at the time of sacrifice as well as 11 months after injection of same activities (in tumor-free mice). However, dendrimer-radioconjugates delivered significant doses to the kidneys and heart, which may ultimately be dose-limiting. Importantly, however, the presented therapeutic benefit and prolonged survival were observed at administered activities of dendrimer-radioconjugates which were below the MTD and were, separately, shown to not cause long term (after 11 months) renal and/or cardiac toxicities (Figure S8).

The above observation is attributed to the uniquely short range of α-particles in tissue and to the relatively different microdistributions of dendrimer-radioconjugates in the tumors - more uniform due to TAMs infiltration - and in normal organs (less uniform). For α-particle radiation, it is not only the levels of absorbed doses but also the (micro)distributions of the absorbed doses within the tumor and/or normal organs that determine the therapeutic/toxicity effects [35]. Uniform distribution of αRPT increases the probability that *every* cancer cell can be hit by an α-particle. Non-uniform distributions of αRPT usually result in lower efficacy/toxicities [8, 28, 35, 36] because only part(s) of the tumor (and/or tissue) are affected since fewer different cells are being hit. This is true both for tumors and for normal organs. We have previously demonstrated that lower absorbed doses, delivered uniformly in tumors [8, 28, 29, 35], inhibited tumor growth better than higher absorbed doses delivered in a heterogeneous way. Accordingly, in normal organs, spatially heterogeneous α-irradiation, even at significant absorbed doses, may not result in notable toxicities (see Figure S12 in [8], and [36]). These differential patterns of αRPT microdistributions between the tumor and normal organs enabled us to operate at administered activities below the MTD and still observe therapeutic effects while keeping toxicities low [8, 28, 29, 37], although some normal organ absorbed doses were greater than the tumor absorbed doses.

The ability of dendrimers to target reactive microglia/macrophages is not unique to this animal model but has been shown in multiple preclinical models of disease [15]. Additionally, dendrimers, without the radioactivity, have been demonstrated to be safe in humans in completed Phase 1/ 2 trials [16, 38].

In mice, the permeability of the BB<u>T</u>B enabled adequate delivery of dendrimer-radioconjugates following systemic administration. On other preclinical models and/or in humans, if BB<u>T</u>B permeability were to be deemed inadequate for the delivery of effective doses, intraarterial injection could be a clinically relevant alternative. The significance of our approach would not be compromised even in the case of locoregional administration, because it addresses a major challenge of treating tumors in the brain: not how to deliver the therapeutic agents to the tumor (that is affected by the permeability of the BB<u>T</u>B), but how to “spread” the agents within the tumor [4]. Dendrimer-radioconjugates address the latter challenge by employing the intrinsically tumor-infiltrating TAMs [13], which, importantly, are less sensitive to αRPT relative to glioblastoma cells.

## Conclusion

Systemically injected dendrimer-radioconjugates that target glioblastoma-residing tumor-associated macrophages, prolong survival; this is further enhanced by pre-injection of non-lethal doses of standard-of-care chemotherapy, without notable toxicities.

## Acknowledgements

The authors thank Dr. George Sgouros, for support with the dosimetry calculations, and Dr. Laura Ensign, for support with the animal studies. This work was supported by the Biomedical Research Award from Hartwell Foundation.

## KEY POINTS

### QUESTION

Can the efficacy of αRPT against glioblastoma be improved?

### PERTINENT FINDINGS

Systemically injected dendrimer-radioconjugates that “hitchhike” glioblastoma-residing tumor-associated macrophages, which are inherently tumor-infiltrating, prolong survival. This is further enhanced by pre-injection of non-lethal doses of standard-of-care chemotherapy, without notable toxicities.

### IMPLICATIONS FOR PATIENT CARE

Augmenting standard-of-care chemotherapy, such as temozolomide, with low doses of α-particle radiotherapy delivered by dendrimer-radioconjugates, may safely prolong survival of patients, who have only a few months to live and limited treatment options.

