## Supporting Information for "Glioblastoma Treatment by Systemic Actinium-225 α-particle Dendrimer-radioconjugates is Improved by Chemotherapy"

##### Materials

Ethylenediaminetetraacetic acid (EDTA) was purchased from Fisher Scientific (Pittsburgh, PA, USA), and Phosphate Buffered Saline (PBS) was purchased from Sigma-Aldrich (Atlanta, GA, USA). The 3kDa MWCO Amicon Ultra-0.5 Centrifugal Filter Unit (Cat. No. UFC5003) was purchased from Milli Pore Sigma (Saint Louis, MO, USA). Matrigel™, Trypsin and the ultra-low adhesion U-shaped 96-well plates (Cat. No.: 7007) were purchased from Corning (Corning, NY, USA), the Dulbecco's Modified Eagle Medium (DMEM) and Roswell Park Memorial Institute (RPMI) medium were from ATCC (Manassas, VA, USA), the Fetal Bovine Serum (FBS) was from Omega Scientific (Tarzana, CA, USA) and penicillin-streptomycin and SNARF-4F were from ThermoFisher Scientific (Waltham, MA, USA). Cyanine 5 (Cy5) was from GE Healthcare Life Science (Pittsburgh, PA). The recombinant murine IL-4 (Cat. No. 214-14) was purchased from Peprotech (Cranbury, NJ, USA). Temozolomide was purchased from Tokyo Chemical Industry (Portland, OR, USA). The 3-(4,5-dimethylthiazol-2-yl)-2,5-diphenyltetrazolium bromide (MTT) assay kit was purchased from Promega (Madison, WI, USA), Chelex® resin from Bio-Rad (Hercules, CA, USA), syringe filters (0.22µm, Cat No. 76479-024) from VWR (Radnor, PA, USA).

#### Methods

*Radiolabeling of dendrimers*

To radiolabel dendrimers with [$^{225}$Ac]Ac (or [$^{111}$In]In), DOTA-dendrimer (or DTPA-dendrimer) was suspended in 500µL of Tris-HCl buffer, pH 9.0 (or acetate buffer, pH 4.5) in Chelex-ed water. [$^{225}$Ac]$AcCl_3$ (or [$^{111}$In]$InCl_3$) dissolved in 0.2M HCl was added, and the reaction mixture was incubated at 37°C for one hour. Dendrimers were then purified using a 3kDa MWCO centrifugation filter, washing 3 times at 12,000 RCF for 15 mins each, with PBS at 1mM EDTA, pH 7.4. The radiolabeling efficiency was calculated as the ratio of the measured activity in the dendrimer suspension after the final wash divided by the activity before the first wash. Radiochemical purity was evaluated using iTLC with 10mM EDTA in water as the mobile phase [1]. The specific activity of dendrimers was measured by counting, for a certain mass of dendrimer, the γ-photon emissions of bismuth-213 at 360-480 keV (after reaching secular equilibrium) or of [$^{111}$In]In at 100-400 keV, using a γ-counter (Packard Cobra II Auto-Gamma, Model E5003). The stability of radiolabeling was assessed 24 hours after incubating radiolabeled dendrimers in FBS-supplemented media, at pH 7.4 and 37°C, and by comparing the specific activity of dendrimers before and after purification by centrifugation.

*Clonogenic cell survival assay and measurement of radioactivity associated per cell*

For evaluating the clonogenic survival fractions, cells (in 6-well plates; 500,000 cells per well) were incubated for 6 hours with actinium-225 and/or TMZ, followed by gentle washing, scraping and re-plating into tissue culture dishes at varying cell densities. Once cell colonies were observed (~6 weeks for cancer cells and ~3 weeks for macrophages), dishes were washed with water, and colonies were fixed and stained using 6% (w/v) glutaraldehyde and 0.05% (w/v) Crystal violet, respectively, and were counted. The number of colonies counted for each of the treatment groups was then normalized by the number of colonies from the non-treated group to obtain the survival fraction, while accounting for the plating efficiency [2].

*Spheroids*

Spheroids were formed by plating 1,000 GL261 cells per well onto ultra-low adhesion U-shaped 96-well plates (in 1.8% v/v Matrigel™) following centrifugation at 1,023 RCF for 10 minutes. Spheroids were then let grow until they reached 400µm in diameter.

To evaluate the spatiotemporal microdistributions within GL261 spheroids of dendrimers (and/or of free TMZ), spheroids were incubated with Cy5-dendrimers (ex/em: 651/670 nm) (and/or CFDA-SE, employed as surrogate of free TMZ; ex/em: 492/517 nm) at a final concentration of 10µg/mL (and/or 10µM) for a total of 3 hours (and/or 15 mins), during which, spheroids were sampled at different time points ("uptake" phase), and were then transferred to fresh media and were sampled during the "clearance" phase. Sampled spheroids were frozen in Cryochrome™ gel over dry ice. The cryomolds were sliced on a HM550 cryotome at 20µm thickness, and the equatorial spheroid slices were imaged on the Zeiss LSM-780 Laser Scanning Confocal Microscope (Zeiss, White Plains, NJ, USA). The radial distributions of the fluorescence intensities of Cy5-dendrimers (and/or of CFDA-SE) were evaluated by applying an erosion code, as described before [3]. Three slices (one per spheroid) were analyzed for each time point. Fluorescence intensities were converted to concentrations using a calibration curve constructed using serial dilutions of entities in a 20μm-pathlength cuvette imaged by the same microscope. Spheroids that were not incubated with fluorophores were used as background.

*Animal study – MRI imaging and confirmation of brain tumors*

To confirm the presence of tumor and to then randomly assign mice to a treatment condition/control group, mouse brain images were acquired on a multi-nuclear BioSpec 70/30 PET-MR 7T scanner (Bruker Biospin MRI Inc., Billerica, MA, USA). The mice were anesthetized under ~1.5-2% isoflurane. The mouse head was imaged in coronal orientation using a T2-weighted fast spin-echo rapid imaging with refocused echo (RARE) sequence (repetition time, 3 s; echo time, 30 ms; RARE factor, 8; number of average, 4; matrix size, 200 x 200; field of view, 2.4 × 2.4 cm; slice thickness, 0.5 mm; in-plane resolution, 120 × 120 μm).

**Table S1**. Tabulated values of decay-corrected biodistributions of the systemically administered 100μL of 740kBq [$^{111}$In]In-DTPA-dendrimers on C57BL/6 mice with intracranial GL261 glioblastoma tumors. Error bars correspond to standard deviations of n=3 mice per condition per time point.

| %IA [$^{111}$In]In-DTPA-dendrimer/g | | | | | | | | |
|---|---|---|---|---|---|---|---|---|
| Time (hours) | 0.25 | 1 | 4 | 8 | 24 | 48 | 72 | 96 |
| Liver | 0.43 ± 0.06 | 1.07 ± 0.05 | 1.50 ± 0.26 | 1.29 ± 0.20 | 0.92 ± 0.12 | 0.36 ± 0.02 | 0.29 ± 0.01 | 0.22 ± 0.03 |
| Spleen | 0.02 ± 0.00 | 0.02 ± 0.00 | 0.02 ± 0.00 | 0.01 ± 0.00 | 0.01 ± 0.00 | 0.01 ± 0.00 | 0.01 ± 0.00 | 0.01 ± 0.00 |
| Kidney | 0.87 ± 0.09 | 2.37 ± 0.52 | 2.78 ± 0.22 | 2.49 ± 0.24 | 2.25 ± 0.28 | 1.04 ± 0.11 | 0.66 ± 0.01 | 0.41 ± 0.06 |
| Lung | 0.03 ± 0.00 | 0.04 ± 0.01 | 0.03 ± 0.01 | 0.05 ± 0.01 | 0.04 ± 0.01 | 0.02 ± 0.01 | 0.02 ± 0.00 | 0.01 ± 0.00 |
| Heart | 2.10 ± 0.59 | 1.47 ± 0.42 | 0.52 ± 0.12 | 0.41 ± 0.11 | 0.36 ± 0.03 | 0.28 ± 0.04 | 0.24 ± 0.05 | 0.16 ± 0.02 |
| Brain | 0.07 ± 0.01 | 0.06 ± 0.02 | 0.09 ± 0.03 | 0.08 ± 0.02 | 0.07 ± 0.01 | 0.04 ± 0.01 | 0.04 ± 0.01 | 0.02 ± 0.00 |
| Tumor | 0.03 ± 0.01 | 0.06 ± 0.00 | 0.10 ± 0.01 | 0.12 ± 0.01 | 0.13 ± 0.01 | 0.12 ± 0.01 | 0.09 ± 0.01 | 0.06 ± 0.01 |
| Blood | 44.5 ± 8.39 | 12.2 ± 1.12 | 8.14 ± 0.70 | 7.43 ± 0.57 | 6.27 ± 1.09 | 3.53 ± 0.36 | 2.26 ± 0.30 | 1.74 ± 0.09 |
| Stomach | 0.04 ± 0.01 | 0.05 ± 0.01 | 0.06 ± 0.01 | 0.04 ± 0.00 | 0.04 ± 0.01 | 0.05 ± 0.02 | 0.05 ± 0.01 | 0.04 ± 0.01 |
| Intestine | 0.03 ± 0.01 | 0.05 ± 0.02 | 0.07 ± 0.01 | 0.05 ± 0.01 | 0.06 ± 0.01 | 0.08 ± 0.02 | 0.08 ± 0.01 | 0.06 ± 0.01 |

**Table S2**. Tabulated values of decay-corrected biodistributions of the systemically administered 100μL of 740kBq [$^{111}$In]In-DTPA-dendrimers - injected 24 hours after administration of TMZ (80 mg/Kg) - on C57BL/6 mice with intracranial GL261 glioblastoma tumors. Error bars correspond to standard deviations of n=3 mice per condition per time point.

| %IA [$^{111}$In]In-DTPA-dendrimer/g | | | | | | | | |
|---|---|---|---|---|---|---|---|---|
| Time (hours) | 0.25 | 1 | 4 | 8 | 24 | 48 | 72 | 96 |
| Liver | 0.40 ± 0.06 | 1.11 ± 0.16 | 1.70 ± 0.11 | 1.42 ± 0.22 | 0.97 ± 0.03 | 0.39 ± 0.04 | 0.32 ± 0.02 | 0.23 ± 0.03 |
| Spleen | 0.02 ± 0.00 | 0.02 ± 0.00 | 0.01 ± 0.00 | 0.01 ± 0.00 | 0.01 ± 0.00 | 0.01 ± 0.00 | 0.01 ± 0.00 | 0.01 ± 0.00 |
| Kidney | 0.89 ± 0.02 | 2.42 ± 0.35 | 2.75 ± 0.17 | 2.34 ± 0.41 | 2.29 ± 0.18 | 1.02 ± 0.20 | 0.60 ± 0.01 | 0.38 ± 0.07 |
| Lung | 0.03 ± 0.001 | 0.04 ± 0.01 | 0.04 ± 0.00 | 0.04 ± 0.01 | 0.05 ± 0.01 | 0.02 ± 0.00 | 0.02 ± 0.00 | 0.01 ± 0.00 |
| Heart | 1.90 ± 0.24 | 1.23 ± 0.14 | 0.51 ± 0.12 | 0.40 ± 0.07 | 0.36 ± 0.04 | 0.28 ± 0.07 | 0.23 ± 0.04 | 0.20 ± 0.01 |
| Brain | 0.06 ± 0.00 | 0.07 ± 0.01 | 0.09 ± 0.01 | 0.08 ± 0.01 | 0.06 ± 0.01 | 0.05 ± 0.01 | 0.04 ± 0.01 | 0.04 ± 0.01 |
| Tumor | 0.04 ± 0.01 | 0.07 ± 0.01 | 0.13 ± 0.01 | 0.14 ± 0.01 | 0.16 ± 0.01 | 0.15 ± 0.01 | 0.13 ± 0.02 | 0.08 ± 0.00 |
| Blood | 43.3 ± 5.66 | 10.8 ± 2.02 | 8.67 ± 0.83 | 7.32 ± 0.47 | 6.24 ± 0.55 | 3.83 ± 0.39 | 2.53 ± 0.09 | 1.64 ± 0.05 |
| Stomach | 0.04 ± 0.00 | 0.05 ± 0.01 | 0.05 ± 0.01 | 0.05 ± 0.00 | 0.05 ± 0.01 | 0.05 ± 0.00 | 0.05 ± 0.00 | 0.05 ± 0.00 |
| Intestine | 0.04 ± 0.01 | 0.04 ± 0.01 | 0.04 ± 0.01 | 0.07 ± 0.03 | 0.06 ± 0.01 | 0.08 ± 0.01 | 0.07 ± 0.01 | 0.06 ± 0.01 |

**Table S3**: Dosimetry for systemically injected [$^{111}$In]In-DTPA-dendrimer in (A) the absence of Temozolomide (TMZ), and (B) following the intraperitonial injection of TMZ, 80mg/Kg mouse, 24 hours earlier. Calculated for **22.2kBq** [$^{225}$Ac]Ac-DOTA-dendrimer per 20g mouse using RAPID Dosimetry (Baltimore, MD). Reported are the mean values and standard deviations of dose evaluated for n=3 mice per time point. ** p-value<0.01.

| **Organ** | **Absorbed Dose of Dendrimers (Gy)** | |
|---|---|---|
| | **Without TMZ** | **With I.P. pre-injection of TMZ** |
| Tumor | 0.18 ± 0.01** | 0.24 ± 0.01** |
| Heart | 0.69 ± 0.03 | 0.69 ± 0.02 |
| Lungs | 0.00 ± 0.00 | 0.00 ± 0.00 |
| Liver | 0.16 ± 0.01 | 0.17 ± 0.01 |
| Spleen | 0.02 ± 0.01 | 0.02 ± 0.01 |
| Kidneys | 1.58 ± 0.05 | 1.40 ± 0.05 |
| Brain | 0.08 ± 0.00 | 0.10 ± 0.01 |
| Intestine | 0.00 ± 0.00 | 0.00 ± 0.00 |
| Stomach | 0.00 ± 0.00 | 0.00 ± 0.00 |

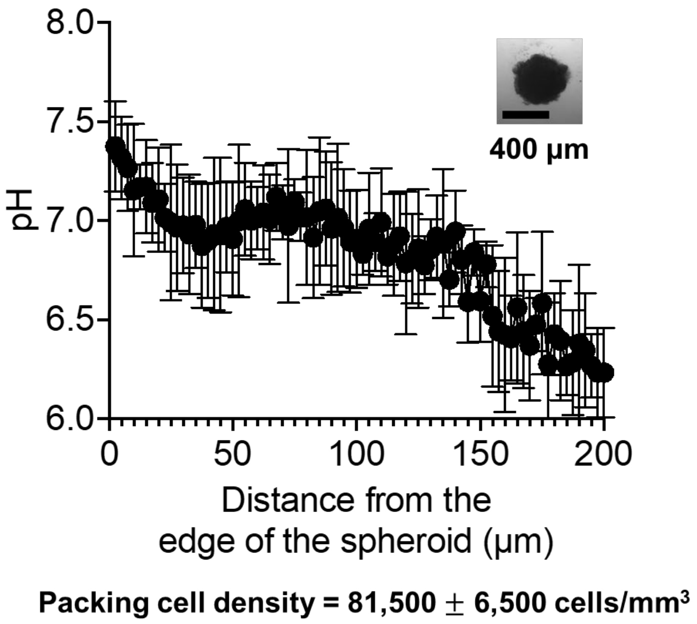


**SUPPLEMENTAL FIGURE 1**. **Interstitial pHe profile in GL-261 spheroids**. Error bars correspond to the standard deviation of repeated measurements (n=6 spheroids, n=3 independent runs).

**Method**

For the measurement of the interstitial $pH_e$ gradient, spheroids with diameters of 400 ± 20 µm, were incubated for 12 hours with 200 µM SNARF-4F, a membrane-impermeant pH sensitive indicator (ex: 488 nm/ em: 580 nm and 640 nm) whose ratio of fluorescence intensities at the two emission wavelengths depend on pH. After incubation, the spheroids were transferred to wells containing fresh media for imaging using a Zeiss LSM 780 Laser Scanning Confocal Microscope, as previously described [4]. An in-house erosion algorithm was applied to the images of the equatorial optical slices to calculate the radial average intensities at each emission channel and their ratios, and to generate the radial pHe maps of spheroids using a calibration curve established by imaging free SNARF- 4F (50 µM) in wells containing media of known pH in the range of interest (7.4−6.0) [5].

The cell density of 400µm-in-diameter GL261 spheroids was measured by counting cells after dissociating spheroids using trypsin (25µL of 0.05% Trypsin mixed with 25µL of PBS) for 2 hours with repeated mixing every 30 minutes. The number of cells was counted manually on a hemocytometer and the cell density was determined as a ratio of the total number of cells counted divided by the spheroid volume. The experiment was performed in triplicates with 10 spheroids per study.

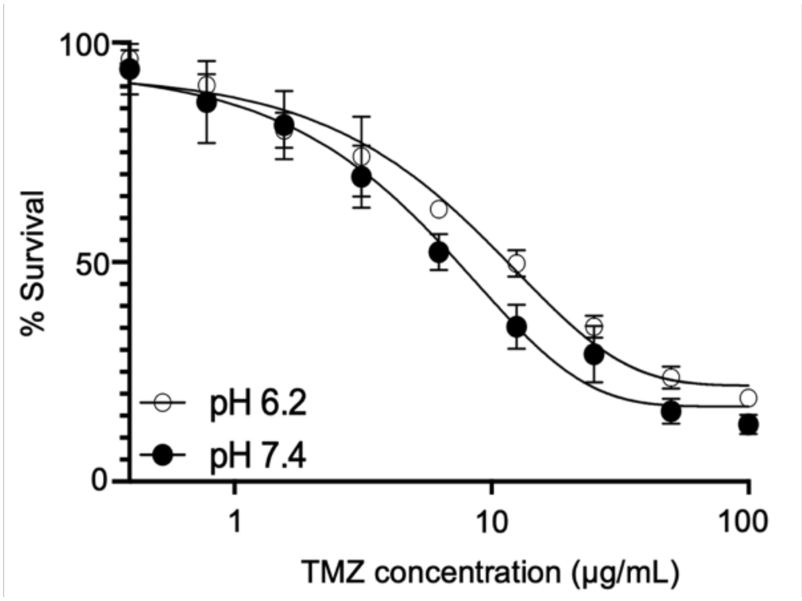


| pH | $IC_{50}$ (µg/mL) |
|---|---|
| 6.2 | 10.27 ± 0.92 |
| 7.4 | 7.53 ± 0.57 |

*

**SUPPLEMENTAL FIGURE 2.** Dose response curves of GL261 glioblastoma cells to increasing concentrations of Temozolomide (TMZ) at pH 6.2 and 7.4. Errors indicate the standard deviations of n = 2 independent runs. * $0.01 < p\text{-value} < 0.05$.

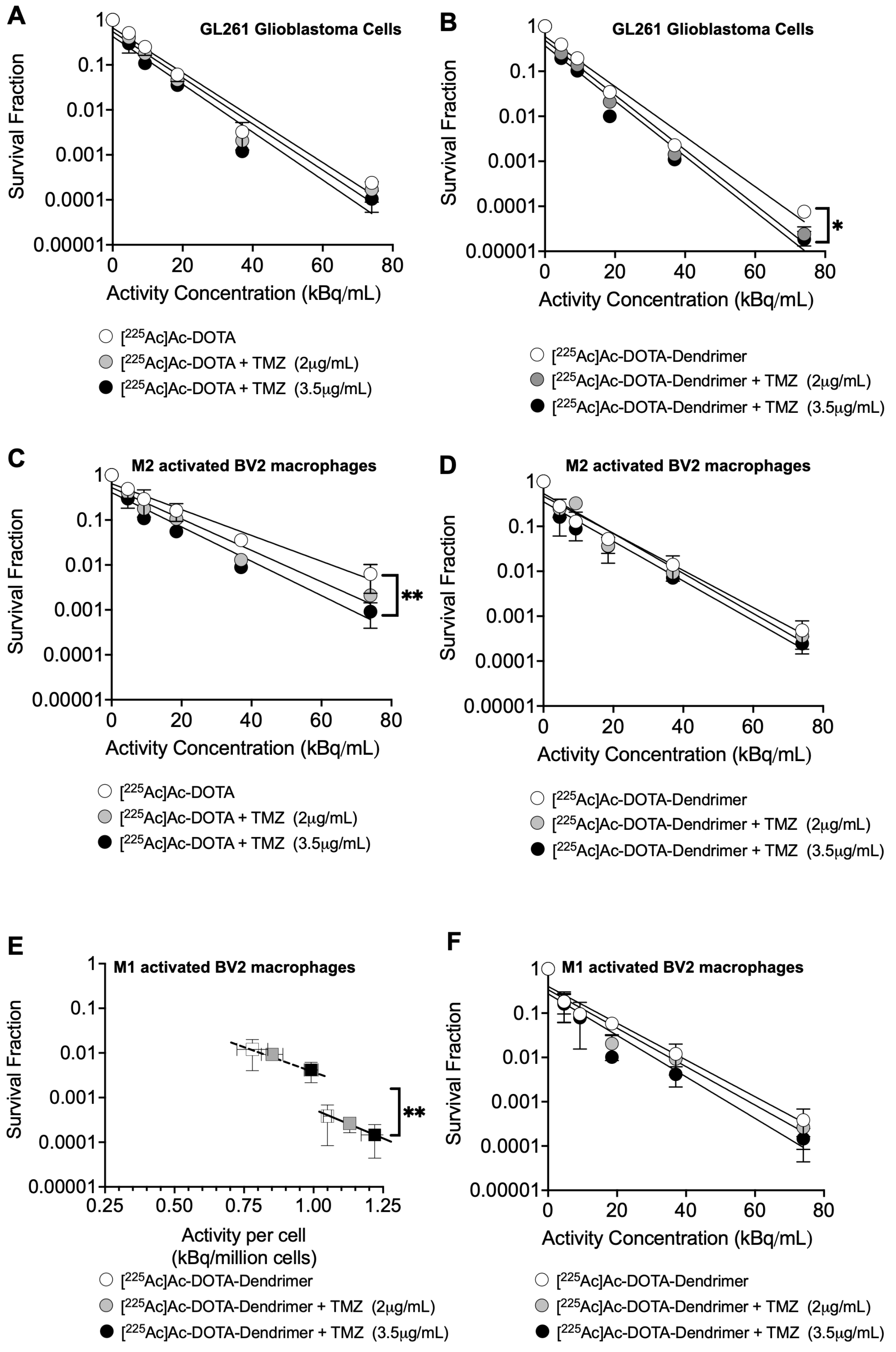


**SUPPLEMENTAL FIGURE 3:** Clonogenic cell survival of GL261 glioblastoma cells (A, B), BV2/M2 macrophages (C, D), and/or BV2/M1 macrophages (F) exposed to [$^{225}$Ac]Ac-DOTA (left hand side) and/or [$^{225}$Ac]Ac-DOTA-dendrimers (right hand side), as a function of activity concentration, in the absence and presence of TMZ at indicated concentrations, for 6 hours at 37°C. (E) Clonogenic cell survival of BV2/M1 macrophages shown in (F)

plotted as a function of the activity associated per cell. The concentration of dendrimer-radioconjugates was kept constant at 10μg/mL at all activity concentrations by adjusting with cold dendrimer. Mean values ± the standard deviation of n=3 independent runs are shown. * indicates $p$-value <0.05; **<0.01.

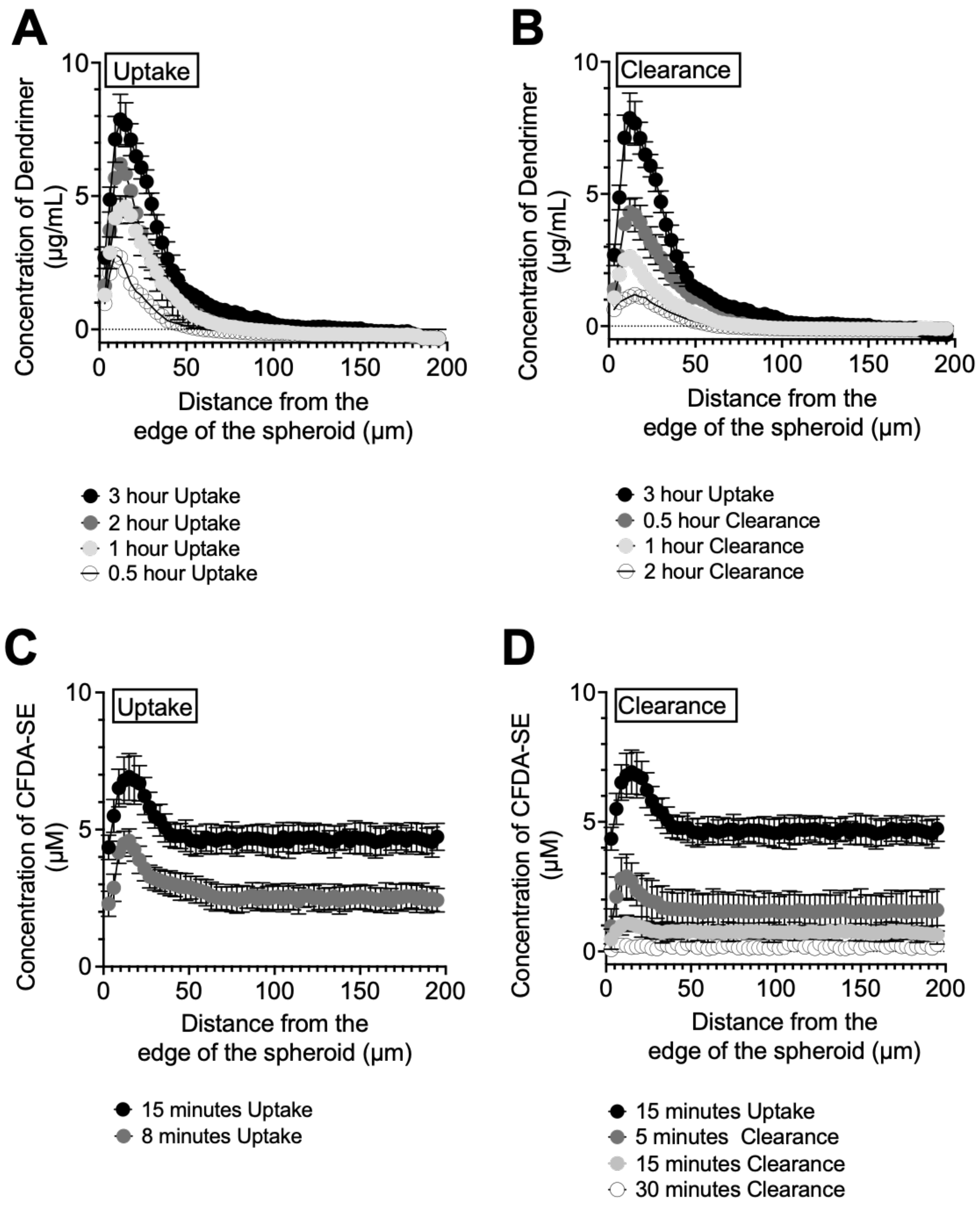


**SUPPLEMENTAL FIGURE 4.** Spatiotemporal profiles of Cy5-dendrimer (A, B) and CFDA-SE (C, D), employed as fluorescent surrogate of temozolomide, in GL261 spheroids. Data points indicate the mean values ± the standard deviations of n=3 different spheroids per time point.

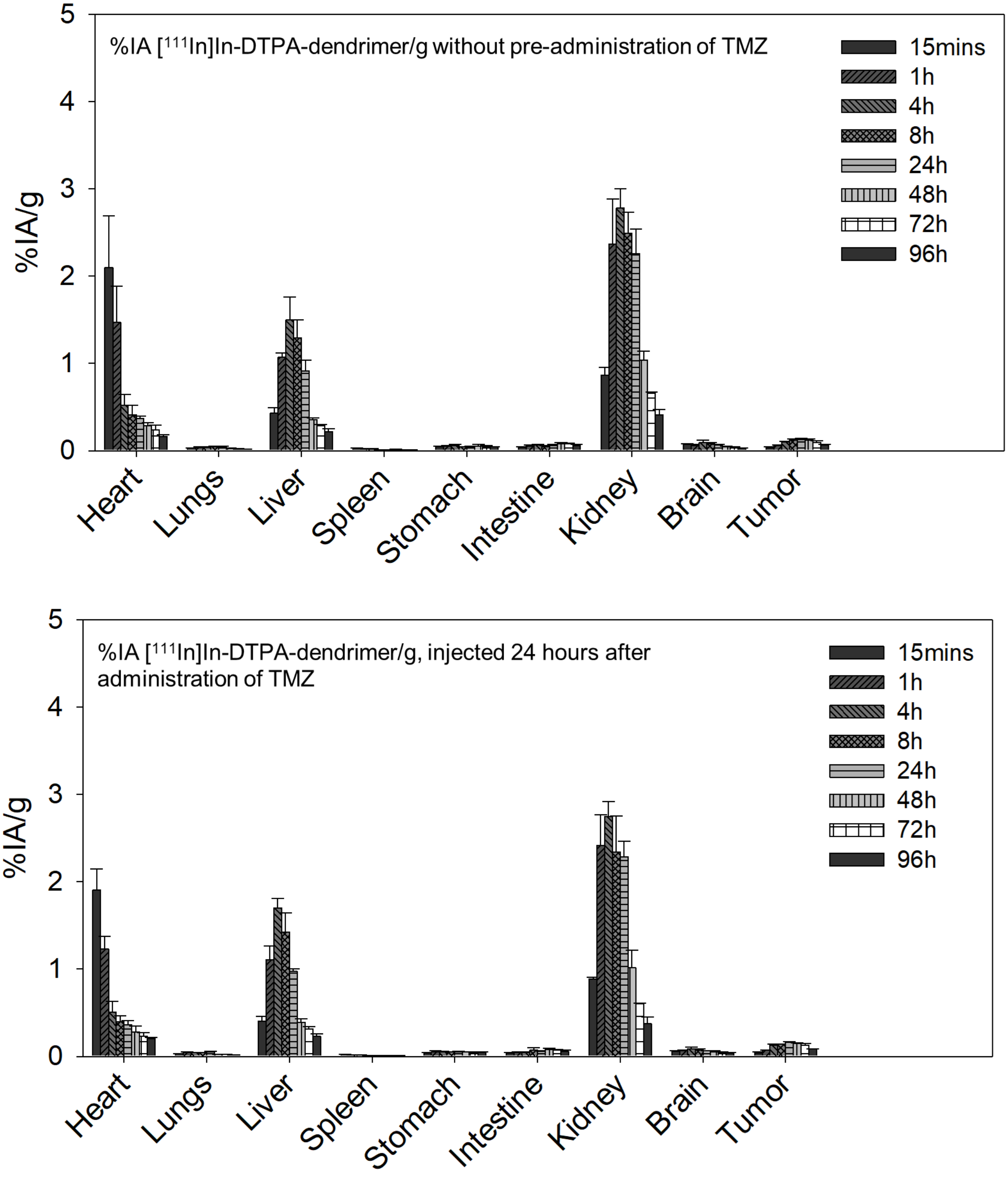


**SUPPLEMENTAL FIGURE 5.** Biodistributions of systemically injected [$^{111}$In]In-DTPA-dendrimer in (A) the absence of Temozolomide (TMZ), and (B) following the intraperitonial injection of TMZ, 80mg/Kg mouse 24 hours earlier, in C57BL/6 mice with intracranial GL261 glioblastoma tumors. Reported are the mean values and standard deviations of dose evaluated for n=3 mice per time point.

### %IA [111In]In-DTPA-dendrimer/g with and without pre-administration of TMZ

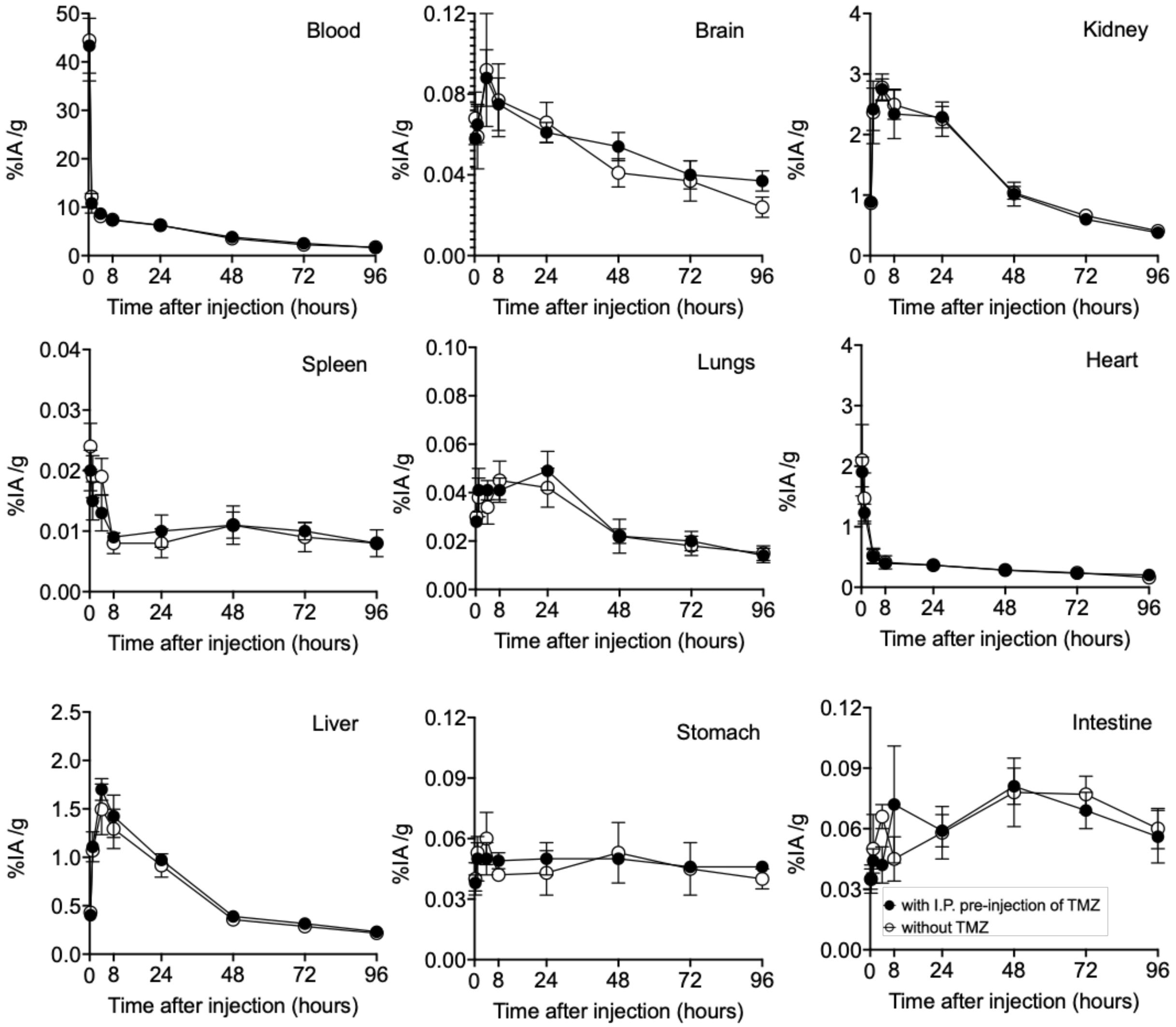


**SUPPLEMENTAL FIGURE 6.** Biodistributions of systemically injected [$^{111}$In]In-DTPA-dendrimer in the absence of Temozolomide (TMZ) (white symbols), and following the intraperitonial injection of TMZ, 80mg/Kg mouse 24 hours earlier (black symbols), in C57BL/6 mice with intracranial GL261 glioblastoma tumors. Reported are the mean values and standard deviations of dose evaluated for n=3 mice per time point.

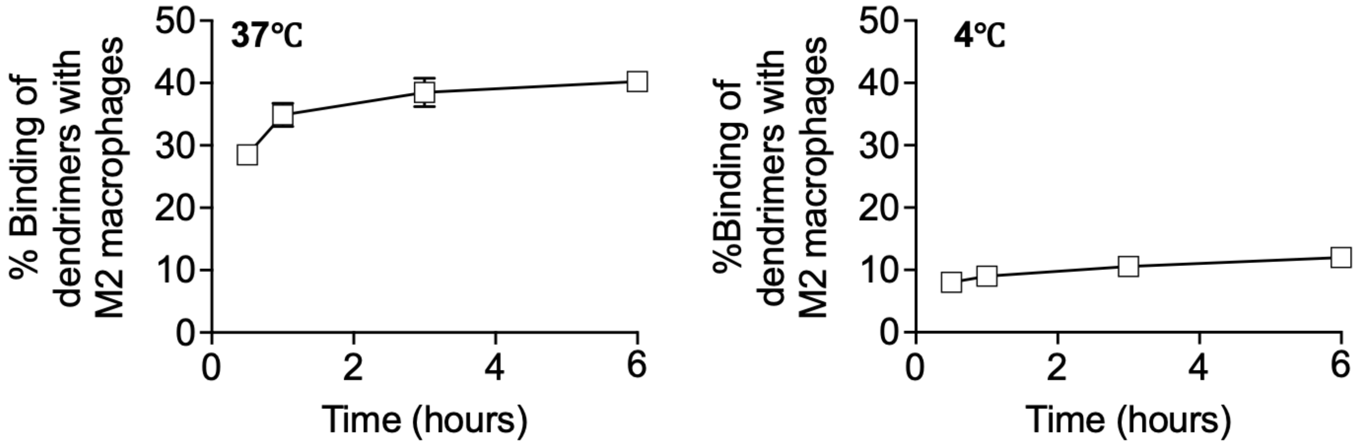


**SUPPLEMENTAL FIGURE 7.** Association of Cy5-dendrimer with M2 activated BV2 macrophages. The difference of the extent of association of dendrimers with macrophages between 37ᵒC and 4ᵒC (after 6 hours of incubation, this difference was 40.23%-11.98% = 28.25%) was primarily attributed to the dendrimer internalization by macrophages that was expected to be highly suppressed at 4ᵒC.

**SUPPLEMENTAL FIGURE 8. Pathology evaluation of long-term (11 months) toxicities** of tumor-free C57BL/6 mice injected systemically with activities **(A) below the MTD** (at **29.6kBq**/ 20g mouse; at which treatment response resulted in significantly prolonged survival, see Figures 6 and 7 of main text), and **(B) above the MTA** (**44.4kBq**/ 20g mouse).

The brain did not raise any long-term (11 months) pathological concerns: neither at 29.6kBq nor at 44.4kBq (on those animals that survived).

Above the MTA, at 44.4kBq, significant renal, splenic and hepatic toxicity was noted. The liver tissue exhibited significant hepatic lipidosis and the kidneys demonstrated tubular atrophy, vacuolization, karyomegaly and slight inflammation and fibrosis. The spleens exhibited iron accumulation, hemosiderin, indicative of past cellular death.

**(A)** Long term (11 month) pathology evaluation of tumor-free C57BL/6 mice injected systemically with 29.6kBq [$^{225}$Ac]Ac-DOTA-dendrimer.

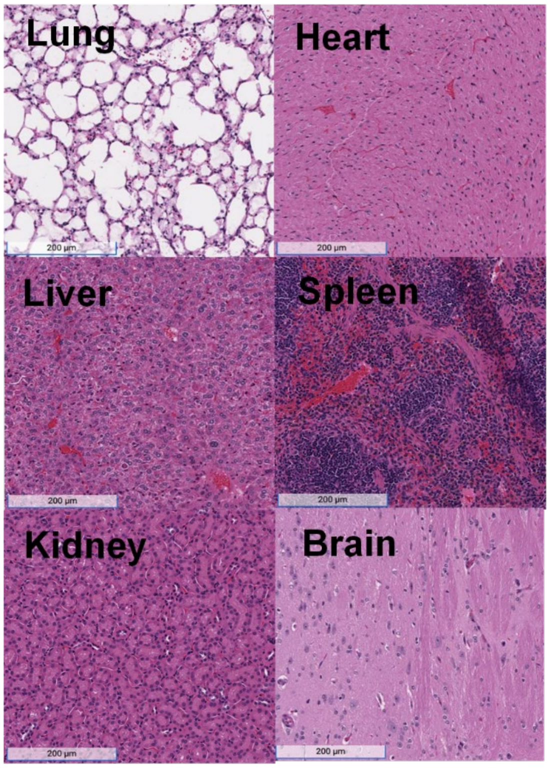


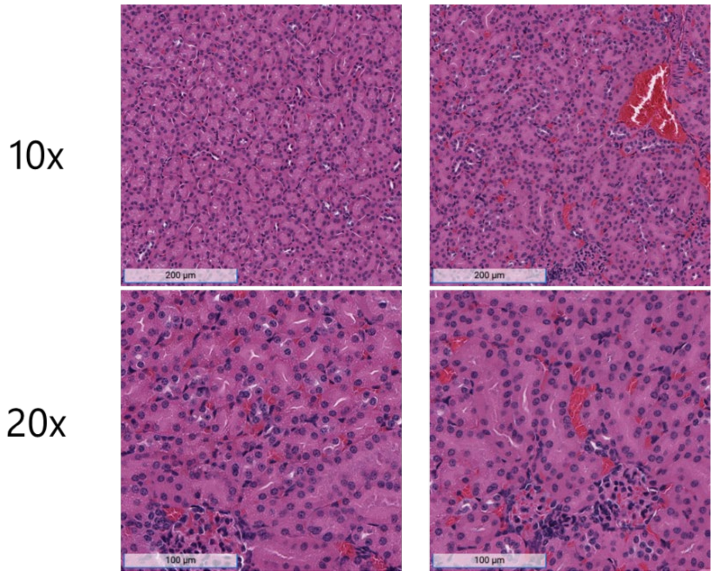


In kidneys, which exhibited significant absorbed doses of the dendrimer-radioconjugate, no noteworthy toxicity findings were observed 11 months after injection of 29.6kBq [$^{225}$Ac]Ac-DOTA-dendrimers.

**(B)** Long term (11 month) pathology evaluation of tumor-free C57BL/6 mice injected systemically with 44.4kBq [$^{225}$Ac]Ac-DOTA-dendrimer, which was above the MTA (defined as the maximum activity at which no deaths were reported).

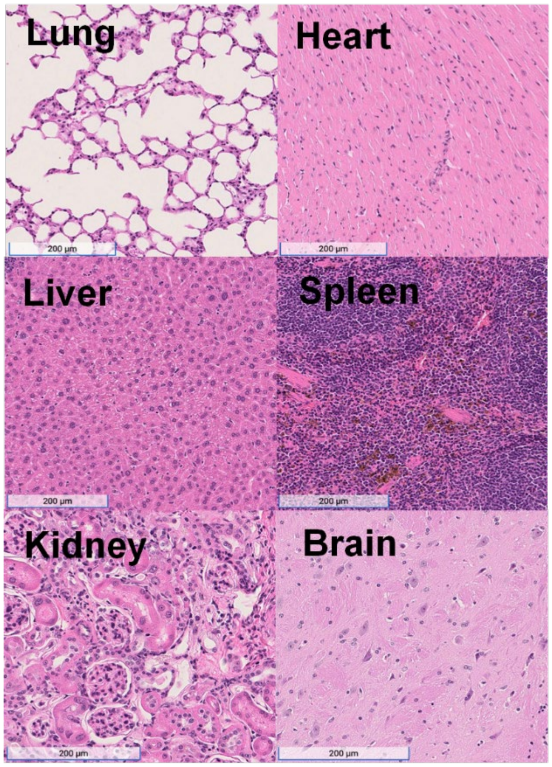


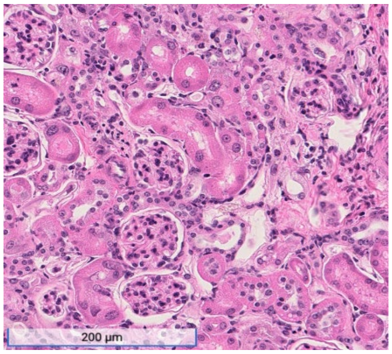


Above the MTD, kidneys exhibited vacuolization, Necrosis, fibrosis, karyomegaly, and inflammation that were observed 11 months after injection of 44.4 kBq [$^{225}$Ac]Ac-DOTA-dendrimers on those mice that survived.

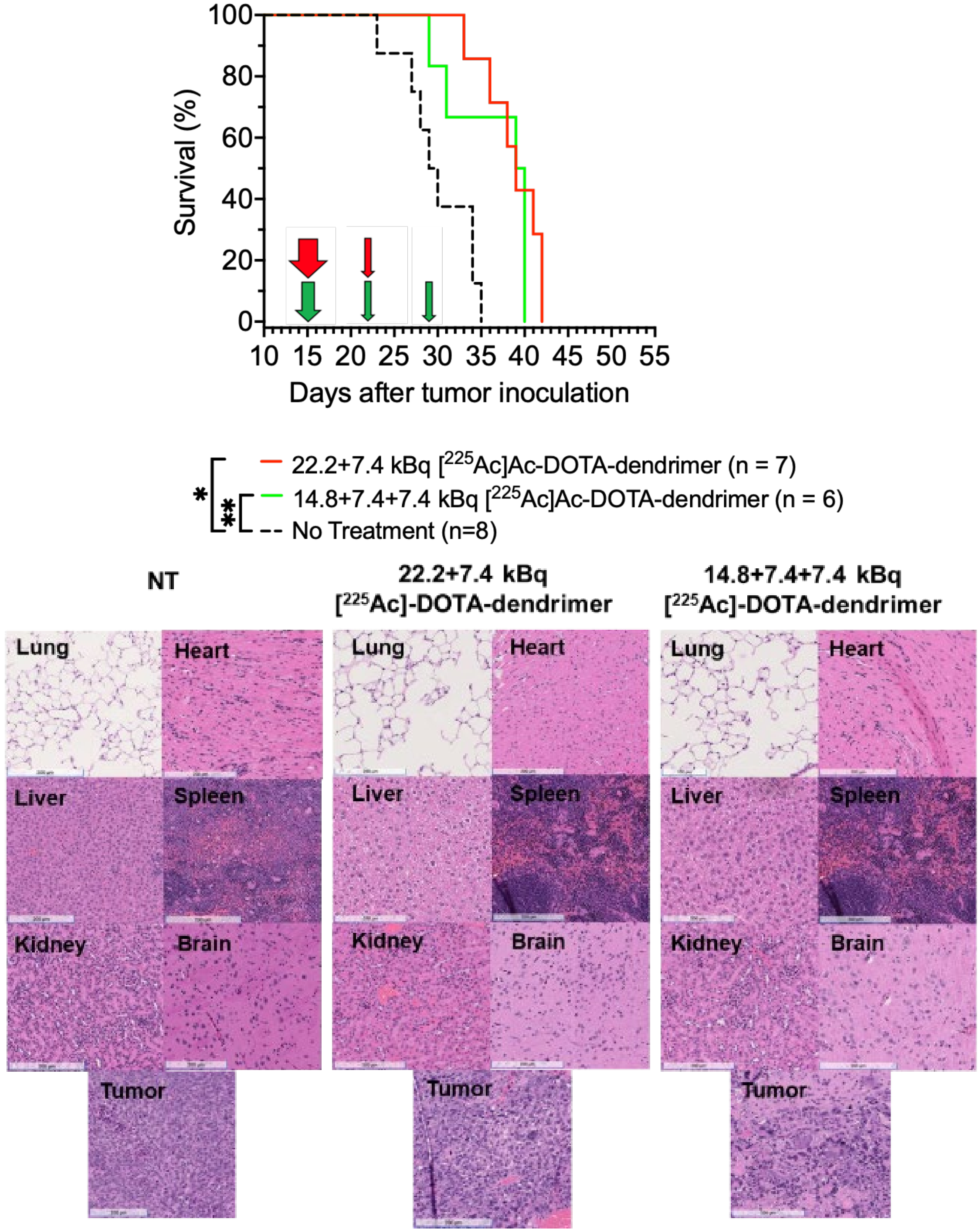


**SUPPLEMENTAL FIGURE 9. Dose fractionation did not affect survival** (top panel), although the time of first death was delayed when a higher fraction of activity was injected first (red line). Vertical arrows indicate treatment scheduling of injected activities shown in the legend. Second panel: H&E stained sections of tissues did not reveal any noteworthy toxicities at the time of sacrifice.

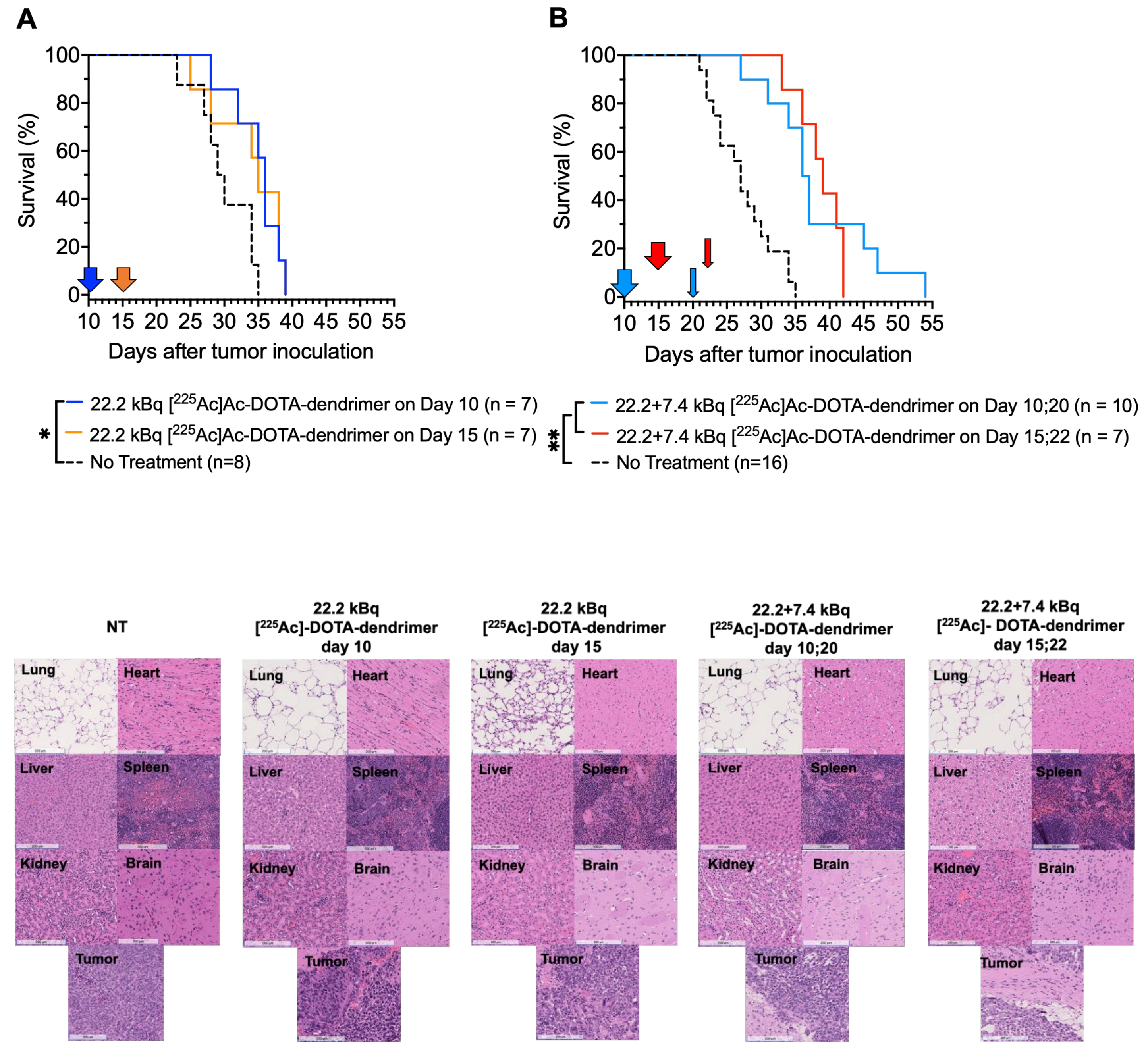


**SUPPLEMENTAL FIGURE 10.** The time of treatment initiation, day 10 (average tumor volume 1.5±0.7 $mm^3$) vs day 15 (9.7±5.7 $mm^3$) after tumor inoculation, did not affect the survival outcome. Animals were treated with (A) 22.2 kBq activity, and (B) 29.6 kBq cumulative activity. Vertical arrows indicate time of treatment. * 0.01<*p*-value <0.05; **<0.01. Bottom panel: H&E stained sections of tissues did not reveal any noteworthy toxicities at the time of sacrifice.

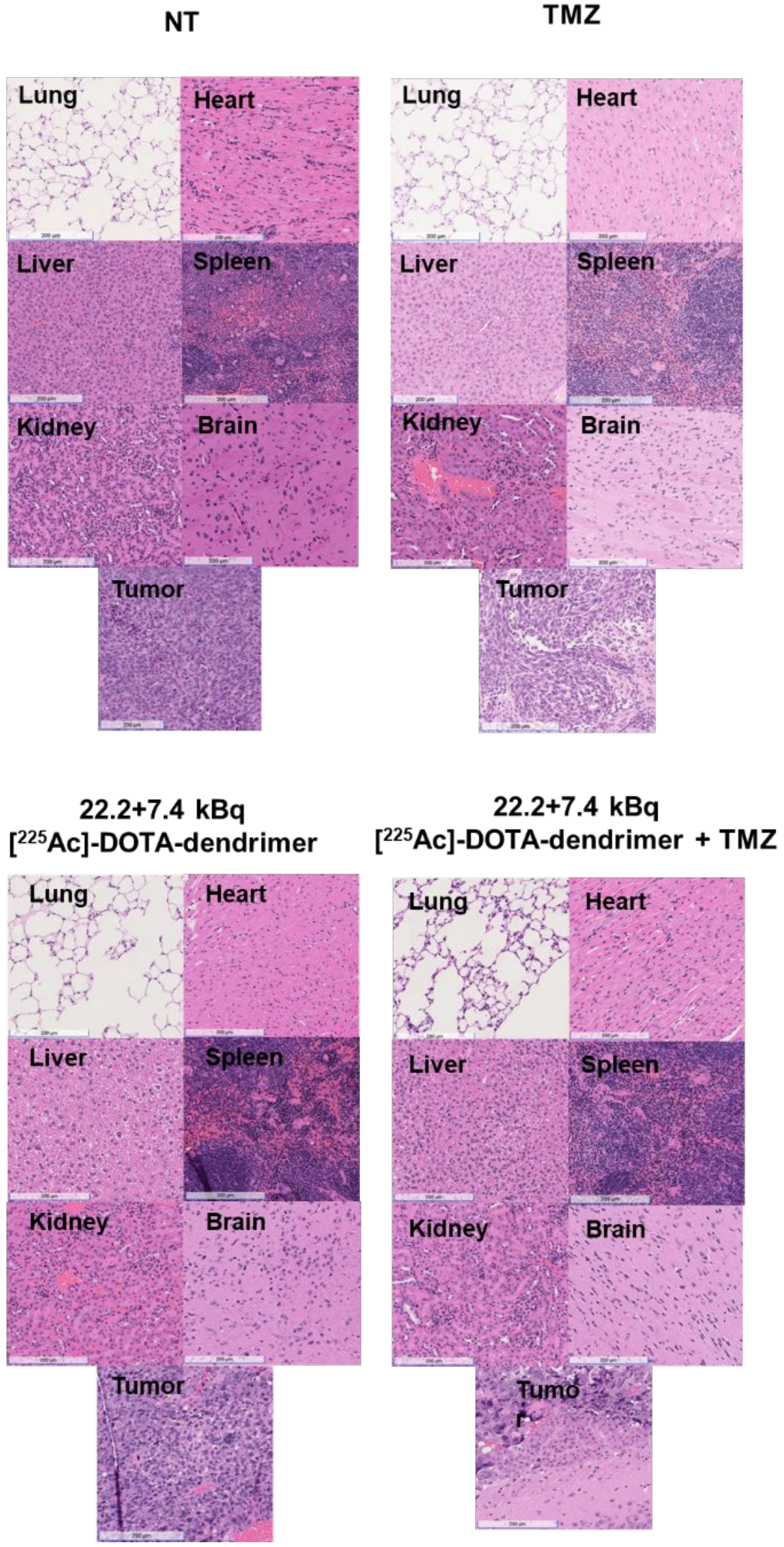


**SUPPLEMENTAL FIGURE 11. H&E-stained sections of tumors and normal organs of mice sacrificed after indicated treatment (as shown in FIGURE 7).** No significant renal, hepatic or splenic toxicity was observed. In the chemotherapy-only cohort, mild hepatic lipidosis was observed. However, this was not exhibited by the cohort treated with the combination of αRPT and chemotherapy. Scale bar=200μm.

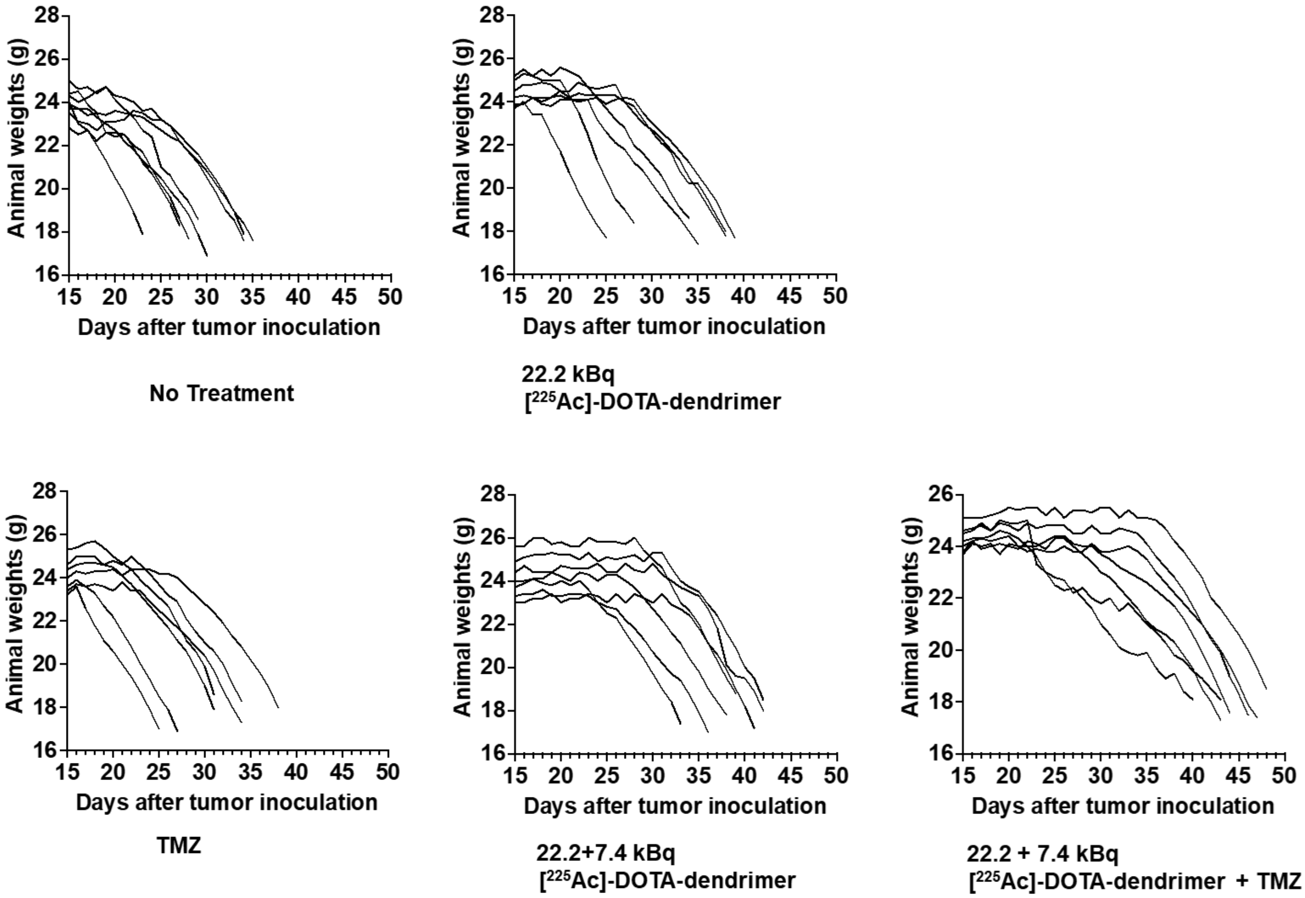


**SUPPLEMENTAL FIGURE 12.** Treatment studies: weight of individual animals during the tumor growth control studies shown in FIGURES 6 and 7.
